# Piezo mechanosensory channels regulate centrosome integrity

**DOI:** 10.1101/2022.04.12.488050

**Authors:** Liron David, Laurel Martinez, Qiongchao Xi, Xueping Fan, Kameron A. Kooshesh, Ying Zhang, Jagesh V. Shah, Weining Lu, Richard L. Maas, Hao Wu

## Abstract

Piezo1 and 2 are evolutionarily conserved mechanosensory cation channels known to function on the cell surface by responding to external pressure and transducing a mechanically activated Ca^2+^ current. Here we show that both Piezo1 and 2 also exhibit concentrated intracellular localization at centrosomes. Both Piezo1 and 2 loss-of-function and Piezo1 activation by the small molecule Yoda1 produced supernumerary centrosomes due to inappropriate centriole disengagement. Using a centrosome-localized GCaMP Ca^2+^-sensitive reporter, we show that perturbations in Piezo modulate Ca^2+^ local concentration at centrosomes. We designed a photoactivable Yoda1 analog (caged-Yoda1) and revealed that its photoactivation specifically at centrosomes leads to rapid premature centriole disengagement within minutes. We identified the sorting nexin Snx5, which is involved in endocytic uptake and trafficking, as a protein-interactor with the conserved Piezo C-terminal domain, and Snx5 also co-localizes with Piezo1 and 2 at centrosomes. Moreover, inhibition of Polo-like-kinase 1 (PLK1) abolishes Yoda1-induced centriole disengagement. Collectively, these data suggest that Piezo1 and 2 in pericentrosomal endosomes control centrosome integrity, likely by maintaining local Ca^2+^ within a defined range through mechanotransduction of cell intrinsic forces from microtubules.

---

Piezo1 and Piezo2 are mechanosensory cation channels discovered as the primary responders to cell membrane pressure in vertebrates with homologs in other organisms^1^. Because of their Ca^2+^- permeability, Piezo1 and Piezo2 can transduce a mechanically activated (MA) Ca^2+^ current into cells^2–4^ and thus perform a wide range of biological functions from vascular development, blood pressure control^5–8^ and red cell volume control^9–11^, to touch sensation and proprioception^12–15^. Both gain- and loss-of-function *PIEZO1* and *PIEZO2* mutations are associated with severe human diseases^16^. In particular, we and others have shown that *PIEZO2* gain-of-function mutations cause pleiotropic musculoskeletal contracture syndromes, including distal arthrogryposis type 5 (DA5), Gordon syndrome (GS) and Marden-Walker syndrome (MWS), while *PIEZO2* loss-of-function causes distal arthrogryposis with impaired proprioception and touch (DAIPT)^17–21^. In an effort to investigate the function of Piezo2 in muscle development, we discovered a general function of Piezo1 and 2 intracellularly in controlling centriole engagement and centrosome number, suggesting that Piezo proteins may represent an important new class of intracellular mechanotransducers.

## Results

### Piezo1 and 2 localize at centrosomes

To address the role of Piezo proteins in muscle development, we used the undifferentiated C2C12 myoblast cell line for immunofluorescence (IF). Using different antibodies and different fixation methods, we found that Piezo1 and 2 invariably exhibited punctate intracellular distribution with strong foci co-stained with the centrosomal marker γ-Tubulin (Fig. 1a,b, Extended Data Fig. 1a-d). This centrosomal localization of Piezo proteins was further validated in the IMCD3 kidney epithelial cell line^22^, and the Neuro-2A mouse neuroblastoma cell line used in the original identification of Piezo1^1^ (Fig. 1c-d, Extended Data Fig. 1e-f), and by GFP fluorescence in C2C12 cell line stably expressing Piezo1-GFP (Extended Data Fig. 1g-i, Supplementary Video 1). To verify the centrosomal localization *in vivo*, we examined by IF of tissue sections of wildtype (WT) mice and *Piezo2*^-/-^ mice that are less embryonically lethal than *Piezo1*^-/-^ mice^8, 23^. We observed Piezo protein puncta that overlapped with γ-Tubulin-marked centrosomes in WT mice (Extended Data Fig. 2a), and lack of such staining in *Piezo2*^-/-^ littermate (Extended Data Fig. 2b-h). While cell surface, nuclear envelope and endoplasmic reticulum localizations have all been reported for Piezo proteins^24–26^, centrosomal localization has not been recognized previously.

**Fig. 1.**
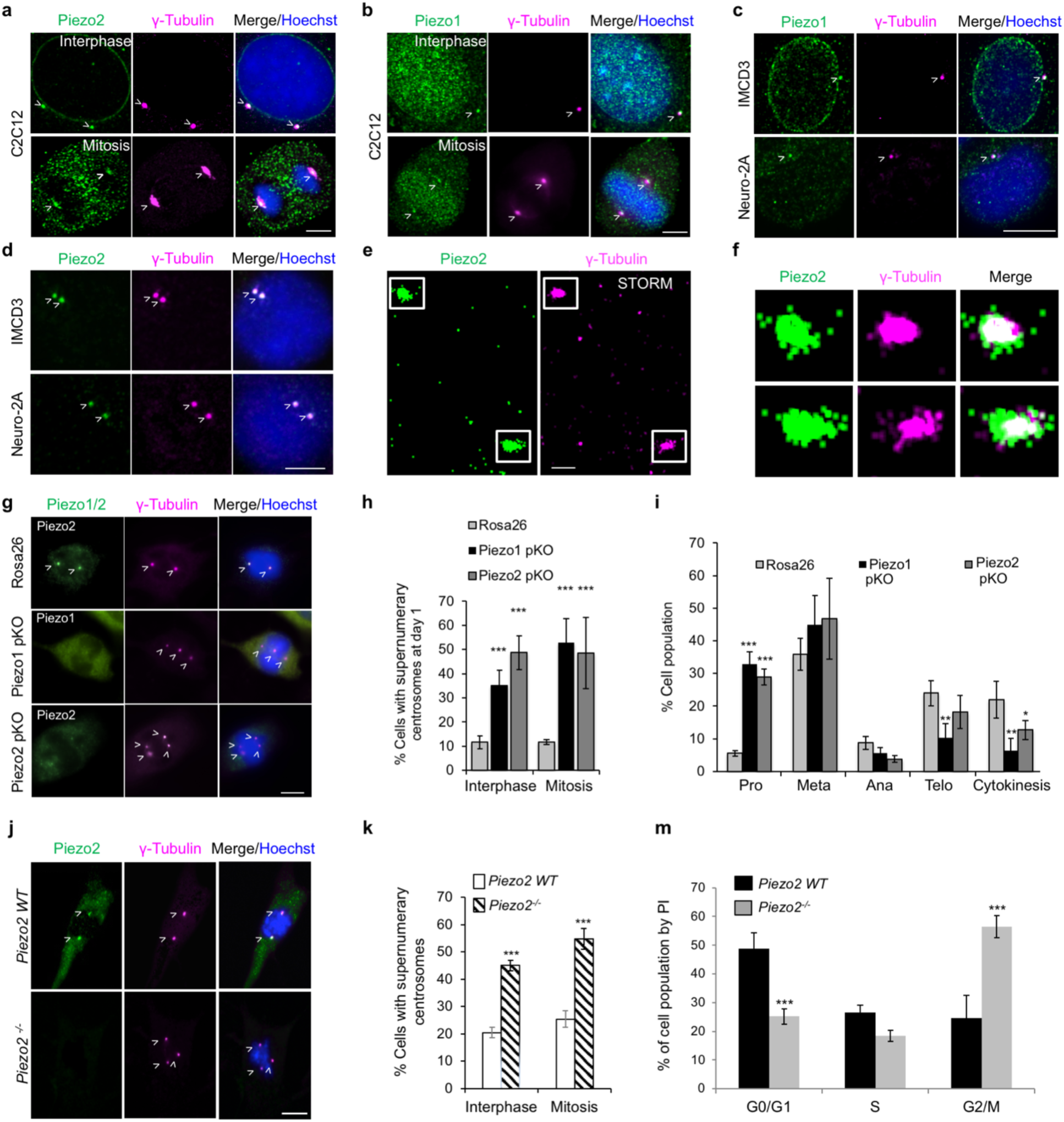
Piezo1/2 localization to centrosomes and the supernumerary centrosome phenotype associated with Piezo1/2 pKO or *Piezo2* ^-/-^. **a, b,** Centrosome localization of Piezo2 (a) and Piezo1 (b) in C2C12 myoblast cells visualized by IF for Piezo1/2 (green), γ-Tubulin (magenta), and DNA (Hoechst dye, blue) in mitotic and interphase cells. **c, d,** Centrosome localization of Piezo2 (d) and Piezo1 (c) in IMCD3 and Neuro-2A cells visualized by IF in interphase cells as in (a-b). **e-f,** STORM imaging performed for fixed unsynchronized C2C12 cells, stained with Piezo2 (green) and γ-Tubulin (magenta). The STORM image (e) showed co-localization of Piezo2 and γ-Tubulin, suggesting Piezo2 localized to the pericentrosomal region, and the insets (f) of the centrosomes are shown with x3 zoom-in. **g,** Rosa26 (off target control), Piezo1 and 2 CRISPR-Cas9 polyclonal KO (pKO) of C2C12 cells at day 1 post selection imaged by IF as in (a-b), showing supernumerary centrosomes in mitotic cells. **h,** Quantitative analysis of supernumerary centrosomes in interphase and mitotic C2C12 pKO cells from the IF experiments in (g). **i,** Quantitative analysis of mitotic cell populations from the IF experiments in (Extended Data fig. 4g). In 3 independent experiments, 190-280 cells were scored for each category based on the mitotic stage. Pro: prometaphase, Meta: metaphase, Ana: anaphase, Telo: telophase, Cyto: cytokinesis, scored as per Methods. **j,** *Piezo2*^-/-^ myoblasts derived from newborn mice imaged by IF as in (a-b), showing supernumerary centrosomes in *Piezo2*^-/-^ myoblasts. **k,** Quantitative analysis of supernumerary centrosomes in interphase and mitotic cells in WT and *Piezo2^-/-^* myoblasts from IF experiments in (j). m, Cell cycle analysis by flow cytometry for WT and *Piezo2^-/-^* myoblasts. All images are maximum intensity Z projections. Centrosomes and centrosome-localized Piezo proteins are marked with white arrowheads. Scale bars are 5 μm for (a-d), 1 μm (e), and 10 μm (g, j). For (h, k), data are represented by mean ± SEM from three independently quantified experiments counting 50-200 cells each. Statistical significance between an experimental group and a control group was assessed by 2-tailed t-test with ***, ** and * denote p < 0.0001, 0.001 and 0.01, respectively.

To more precisely localize Piezo proteins at the centrosome, we performed super-resolution imaging on C2C12 cells co-stained with Piezo2 and γ-Tubulin, using stochastic optical reconstruction microscopy (STORM) (Fig. 1e-f) and instant structured illumination microscopy (iSIM) (Extended Data Fig. 3). Piezo2 co-localized with γ-Tubulin, but was more disperse than γ-Tubulin and even appeared vesicular at the periphery of the foci (Fig. 1f). These data support the pericentrosomal localization of Piezo proteins and are consistent with their association on endosomes (see below).

### Piezo1 or 2 knockout or knockdown induced supernumerary centrosomes and mitotic delay

To investigate the potential function of Piezo proteins on centrosomes, we performed CRISPR-Cas9 polyclonal knockout (pKO) of Piezo1 or Piezo2 in C2C12 cells because selection of KO clones was not possible due to cell death (Extended Data Fig. 4a). Strikingly, we detected by IF imaging many cells with three or more centrosomes, a phenotype known as supernumerary centrosomes^27, 28^, as well as with misaligned spindles (Fig. 1g, Extended Data Fig. 4b, c). Quantification of IF images revealed that 35% to 53% of Piezo pKO C2C12 cells in interphase or mitosis exhibited supernumerary centrosomes, in comparison to 12% for off target Rosa26 pKO (Fig. 1h). Supernumerary centrosomes were also observed in 22 to 45% of C2C12 cells following shRNA knockdown (KD) of Piezo1 or 2 compared to 4 to 10% in control cells receiving a control shRNA (Extended Data Fig. 4d-f). Thus, Piezo1 and 2 are not only localized at centrosomes, but also required for regulation of centrosome number.

To determine if Piezo pKO in C2C12 cells caused mitotic defects, we synchronized these cells at the G2/M border using the cyclin-dependent kinase 1 (CDK1) inhibitor RO-3306 (RO)^29^. After inhibitor washout to release cell cycle arrest, we fixed the cells 45 minutes post-washout. When visualized by IF for both α-Tubulin and γ-Tubulin, C2C12 cells transduced with off target Rosa26 pKO showed predominantly normal dividing cells with bipolar spindles (Extended Data Fig. 4g). Piezo1 and 2 pKO cells at early stages of mitosis (e.g. prometaphase) showed supernumerary centrosomes and misaligned or multipolar spindles; however, if a cell reached late mitotic phase (e.g. cytokinesis), it had predominantly normal centrosomes with bipolar spindles (Extended Data Fig. 4g). Quantitative analysis revealed significant accumulation of Piezo1 and 2 pKO cells in prometaphase compared to off target cells, likely also resulting in less Piezo1 and 2 KO cells in late mitosis (Fig. 1i). These data suggest that the misaligned or multipolar spindle organization in Piezo pKO cells causes mitotic delay

We next sought to determine whether the supernumerary centrosome phenotype associated with Piezo pKO in C2C12 cells also occurred *in vivo.* Primary myoblasts from WT and *Piezo2*^-/-^ littermate newborn mice^14^ were isolated from skeletal muscles and passaged to enrich for myoblasts^30^. Co-IF revealed Piezo2 staining at γ-Tubulin-marked centrosomes and normal numbers of centrosomes in 75 to 80% of WT primary myoblasts (Fig. 1j, k). By contrast, *Piezo2*^-/-^ myoblasts displayed supernumerary centrosomes in 45 to 55% of myoblasts (Fig. 1j, k), resembling those in CRISPR-Cas9 pKO or shRNA KD cells. In addition, cell cycle analysis by flow cytometry revealed a significant accumulation of *Piezo2*^-/-^ cells (56%) in G2/M in comparison to WT cells (25%), with a corresponding decrease in cells in G0/G1 (Fig. 1m).

### Pharmacologic activation and inhibition of Piezo proteins induced supernumerary centrosomes

We employed known pharmacological modulators of Piezo proteins to assess the immediate effects of Piezo activity alteration on centrosomes. Yoda1, a cell-permeable small molecule activator of Piezo1 that induces channel opening in the absence of mechanical force^31, 32^, was used at 10 μM, the mid-point of the previously established dose response curve. C2C12 cells synchronized to the G2/M border by RO for 12 hours were given Yoda1 immediately after RO washout, and analyzed at 30 minutes by IF for α-Tubulin and γ-Tubulin. Approximately 50% of the mitotic cells exhibited supernumerary centrosomes with misaligned or multipolar spindle organization, in comparison to 12% for those that received the DMSO vehicle (Fig. 2a-b, Extended Data Fig. 5a). Co-treatment with 10 μM Yoda1 and 10 μM Dooku1, a modified Yoda1 that binds Piezo1 and antagonizes Yoda1^33^, reduced the percentage of cells with supernumerary centrosomes to 40% (Fig. 2b). With Dooku1 alone, the number of cells with supernumerary centrosomes was 25% (Fig. 2b). At longer time points of 24 and 48 hours post Yoda1 treatment, accumulation of cells in G2 or M phase also became apparent (Extended Data Fig. 5b). Yoda1 treated cells, but not untreated cells, exhibited thin, extended F-actin positive (marked by phalloidin) cell surface protrusions, a phenotype that has been previously observed with Yoda1 treatment^34^ (Fig. 2c, Extended Data Fig. 5c-d).

**Fig. 2.**
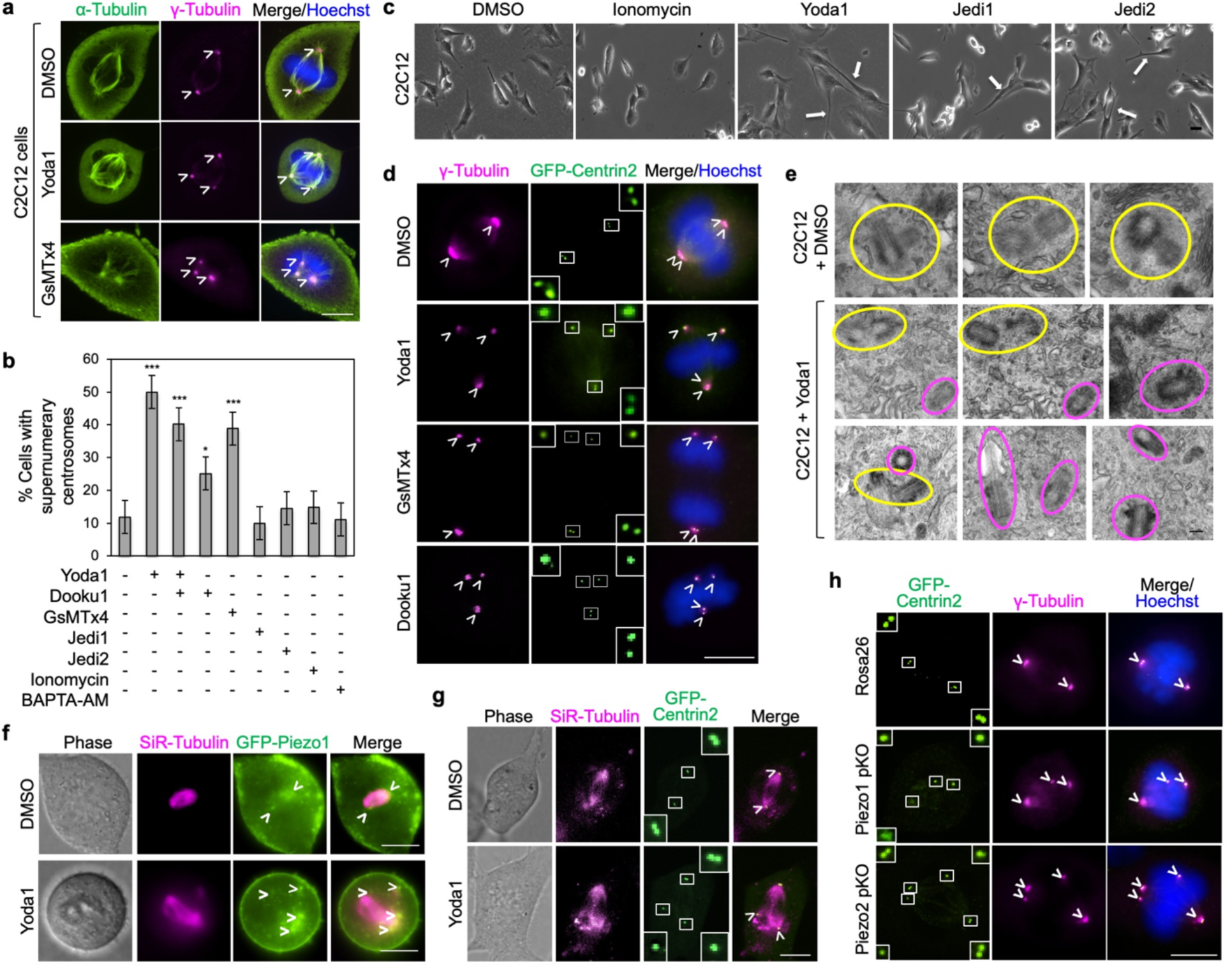
Supernumerary centrosomes upon Piezo pharmacological activation or inactivation and Piezo expression during cell cycle. **a,** C2C12 cells treated for 30 min with DMSO (0.1%, vehicle control), Yoda1 (Piezo1 activator, 10 μM) or GsMTx4 (mechanosensitive channel inhibitor, 5 μM) after RO-3066 release. Cells were imaged by IF for α-Tubulin (green), γ-Tubulin (magenta) and DNA (Hoechst dye, blue). **b,** Quantitative analysis for percentage of C2C12 cells with supernumerary centrosomes from IF images as in (a) following treatment with DMSO (0.1%), Yoda1 (10 μM), Yoda1 plus Dooku1 (10 μM each), Dooku1 (10 μM), GsMTx4 (5 μM), Jedi1 (200 μM), Jedi2 (200 μM), ionomycin (10 μM) or BAPTA-AM (40 μM). **c**, Phase images of C2C12 cells 30 min after treatment with DMSO, ionomycin, Yoda1, Jedi1 and Jedi2 under the same condition as in (b). Yoda1, Jedi1 or Jedi2 treatment, but not DMSO or ionomycin treatment, led to cell surface protrusions due to plasma membrane Piezo1 activation. **d,**C2C12 cells stably expressing GFP-Centrin2 treated for 30 min with DMSO, Yoda1, GsMTX4 or DooKu1 after RO-3066 release, showing supernumerary centrosomes and centriole disengagement. Cells were imaged by IF for γ-Tubulin (magenta), Centrin2 (GFP fluorescence) and DNA (Hoechst dye, blue). Rectangular boxes encircle centrosomes, and their zoom-in views are displayed. **e,** EM gallery of centrosomes, imaged in thin plastic sections of embedded C2C12 cells 1 h after treatment with of DMSO (top) or Yoda1 (bottom). Yellow and magenta circles mark pairs of mother and daughter centrioles, and separated centrioles, respectively. **f, g,** Live fluorescent and phase images of mitotic C2C12 cells stably expressing GFP-Piezo1 (green) (f) or GFP-Centrin2 (g). Cells were treated with DMSO or Yoda1 together with SiR-Tubulin for staining microtubules (magenta) after release of RO-3306-mediated cell cycle synchronization, and imaged 30 min later. Misaligned spindles following Yoda1 treatment are apparent from SiR-Tubulin staining. Rectangular boxes in (g) encircle centrosomes, and their zoom-in views are displayed. **h,** Piezo1 and 2 pKO C2C12 cells stably expressing GFP-Centrin2 after day 1 post-KO selection and RO-3306 synchronization. Cells were washed out and visualized by IF for Centrin2 (GFP fluorescence), γ-Tubulin (magenta) and DNA (Hoechst, blue). All images are maximum intensity Z projections. All scale bars are 10 μm except in (e) which is 200 nm. Data are represented by mean ± SEM from three independently quantified experiments counting 100-250 cells each. Statistical significance was assessed between an experimental group and a control group by 2-tailed t-test with ***, ** and * for p < 0.0001, 0.001 and 0.01, respectively.

To investigate whether the effects of Yoda1 and Dooku1 were due to cell surface or intracellular Piezo1 activation, we used Piezo1 activators Jedi1 and Jedi2, which activate Piezo1 only from the extracellular side and are not cell permeable shown by patching clamping^35^. Treatment of C2C12 cells with either Jedi1 or Jedi2 at 200 µM, a concentration at or above the mid-point of the dose response curve^35^, did not cause an increase in supernumerary centrosomes (Fig. 2b, Extended Data Fig. 5a), but resulted in cell surface protrusions similar to those seen with Yoda1 treatment (Fig. 2c, Extended Data Fig. 5d), supporting that Jedi1 and Jedi2 exerted their expected effect. In addition, we used ionomycin, a Ca^2+^ ionophore that induces intracellular Ca^2+^ influx, and found that ionomycin did not induce supernumerary centrosomes and did not lead to cell surface protrusions (Fig. 2b, c, Extended Data Fig. 5a). These data suggest that aberrant activation of intracellular Piezo1, but not of cell surface Piezo1 or general Ca^2+^ influx, deregulates normal centrosomal number.

We used GsMTx4 peptide, a spider venom toxin that inhibits Piezo1 and 2 as well as other mechanosensitive cation channels^36–39^ to investigate the consequence of Piezo pharmacologic inhibition. Interestingly, similar to Yoda1 treatment, GsMTx4 treatment also significantly increased the number of cells with supernumerary centrosomes to 39% (Fig. 2a-b, Extended Data Fig. 5a). As with lack of centrosomal phenotype when treated with ionomycin, treatment with BAPTA-AM, a cell-permeable Ca^2+^ chelator, also did not lead to supernumerary centrosomes in comparison with untreated cells (Fig. 2b). Thus, both pharmacologic activation of Piezo1 and inhibition of mechanosensitive ion channels including Piezo1 and 2 produced supernumerary centrosomes. While the supernumerary centrosome phenotype with Yoda1 was surprising, the appearance of a similar phenotype with GsMTx4 treatment, presumably reflecting Piezo inhibition, is consistent with the results in Piezo pKO and KD C2C12 cells and in *Piezo2*^-/-^ myoblasts.

### Piezo activation and inhibition induced illegitimate centriole disengagement

To determine whether the supernumerary centrosomes that form upon Piezo activation or inhibition result from centrosome over-duplication, premature centriole disengagement, centrosome fragmentation or some other mechanism^40–42^, we generated a C2C12 cell line that stably expressed GFP-Centrin2. Unlike γ-Tubulin, a component of the pericentriolar material that does not distinguish each centriole, Centrin2 is a component of the centriole per se, and can individually mark mother and daughter centrioles. We added 10 μM Yoda1 to GFP-Centrin2 C2C12 cells, either synchronized at G2/M by RO (Fig. 2d) or unsynchronized (Extended Data Fig. 6a), and analyzed the cells for GFP and by anti-γ-Tubulin IF. Strikingly, upon Piezo1 activation by Yoda1, illegitimate centriole disengagement of one or two centrosomes occurred, resulting in the formation of individual centrioles. Separated centrioles were also identified following either GsMTx4 or Dooku1 treatment (Fig. 2d).

For DMSO control after RO treatment, we did not observe separated centrioles. The low percentage of cells with supernumerary centrosomes in these cells resulted from centriole duplication (Extended Data fig. 6b). The average distance measured between the two disengaged centrioles 30 min post Yoda1 treatment was 4.97 ± 0.85 μm, compared to 0.56 ± 0.03 μm before treatment and 0.54 ± 0.02 μm 30 min post DMSO treatment (Extended Data Fig. 6c). The same centriole disengagement was observed upon Yoda1 treatment when Pericentrin, another component of the pericentriolar material as γ-Tubulin, was used in co-IF to mark the centrosome location (Extended Data Fig. 6d). We also used transmission electron microscopy (TEM) to confirm the distance of disengagement between free centrioles, which was frequently >3.5 μm, compared to ∼200 nm in wild type C2C12 cells (Fig. 2e, Extended Data Fig. 6e).

For distinguishing mother and daughter centrioles, we used Cep164, a mother centriole marker. By imaging mitotic C2C12 cells treated with Yoda1 and performing co-IF for Cep164 and γ-Tubulin, we observed that upon illegitimate centriole disengagement, the daughter centriole preferentially moved away from its original position at one end of the mitotic spindles, while the mother centriole appeared to remain stationary at the end of the mitotic spindle (Extended Data Fig. 6f). In order to visualize the effect on mitotic spindle upon treatment with Yoda1, we imaged Piezo1-GFP C2C12 and GFP-Centrin2 C2C12 cells with the live cell dye SiR-Tubulin to stain microtubules. We identified misaligned spindles following 10 μM Yoda1 stimulation (Fig. 2f, g), as were also observed in Piezo1 or 2 pKO cells (Extended Data Fig. 4c). To extend these findings to other cells, we tested the IMCD3 cell line, which also exhibited Piezo1 and 2 co-localization with centrosomes (Fig. 1c, d). Similar to C2C12 cells, treatment with Yoda1 or GsMTx4 resulted in 40% or 42% of IMCD3 cells, respectively, having supernumerary centrosomes and multipolar spindles 30 min after treatment (Extended Data Fig. 6g, h).

### Piezo knockout induced illegitimate centriole disengagement

Centriole disengagement as a potential mechanism for supernumerary centrosomes in both Yoda1 and GsMTx4-treated C2C12 cells prompted us to consider whether a similar mechanism could account for the supernumerary centrosomes in Piezo1 and 2 pKO cells. We performed CRISPR-Cas9 polyclonal KO of Piezo1 and 2 in C2C12 cells stably expressing GFP-centrin2, and imaged the locations of GFP-Centrin2 and γ-Tubulin. The supernumerary centrosomes in these cells contained either two centrioles, or a single centriole as observed for cells with pharmacologically activated or inhibited Piezo proteins (Fig. 2h, Extended Data Fig. 6i). Thus, centrosome disengagement to form single centrioles occurs in Piezo pKO cells. However, likely due to the longer duration of the KO protocol, centriole replication to form intact centrosomes also occurred^28^.

To test this idea further, we imaged GFP-Centrin2 C2C12 cells after either a 30 min or an extended 24-hour incubation with Yoda1, or after a 30-minute treatment with Yoda1 followed by a 24-hour recovery. Each treatment yielded progressively lower percentages of cells with isolated centrioles and increasing numbers with supernumerary centrosomes containing paired centrioles (Extended Data Fig. 6j, k). These results are also consistent with the idea that free centrioles form in Piezo pKO cells, with some replicating to form intact centrosomes over time. Thus, both Piezo loss- and gain-of-function states appear to produce similar illegitimate centriole disengagement phenotypes, suggesting that a critical range of Piezo activity is required to maintain centrosome integrity likely by mechanotransduction of microtubule forces.

### Rapid illegitimate centriole disengagement upon Piezo1 activation in all cell cycle stages except metaphase

To gain further insight into the kinetics of the centriole dissociation process, we imaged both mitotic and interphase cells in the presence of Yoda1. For imaging mitotic cells, we synchronized GFP-Centrin2 C2C12 cells that also expressed H2B-mCherry to mark chromatin at G2/M with RO. At 15 min, 30 min or 45 min after RO washout, we added Yoda1 or DMSO vehicle, and imaged cells at early, middle or late stage of mitosis, respectively (Fig. 3a, b, Extended Data Fig. 7a, b and Supplementary Video 2-5). For imaging interphase cells, we used unsynchronized cells in G1 and G2, added Yoda1 and recorded centriole disengagement events (Fig. 3c, Supplementary Video 6-7). With DMSO vehicle, intact centrosomes migrated to the spindle poles and the centrioles maintained tight coupling throughout the imaging period (Extended Data Fig. 7a, Supplementary Video 2). With Yoda1 treatment, we observed rapid centriole disengagement within a few minutes for cells at early mitotic stage (prophase), resulting in misaligned spindles and lagging chromatin marked by H2B-mCherry (Fig. 3a, Supplementary Video 3). Rapid centriole disengagement was also observed for cells at late mitotic stage (cytokinesis) or interphase at G1 and G2 (Fig. 3b, c, Supplementary Video 5-7).

**Fig. 3.**
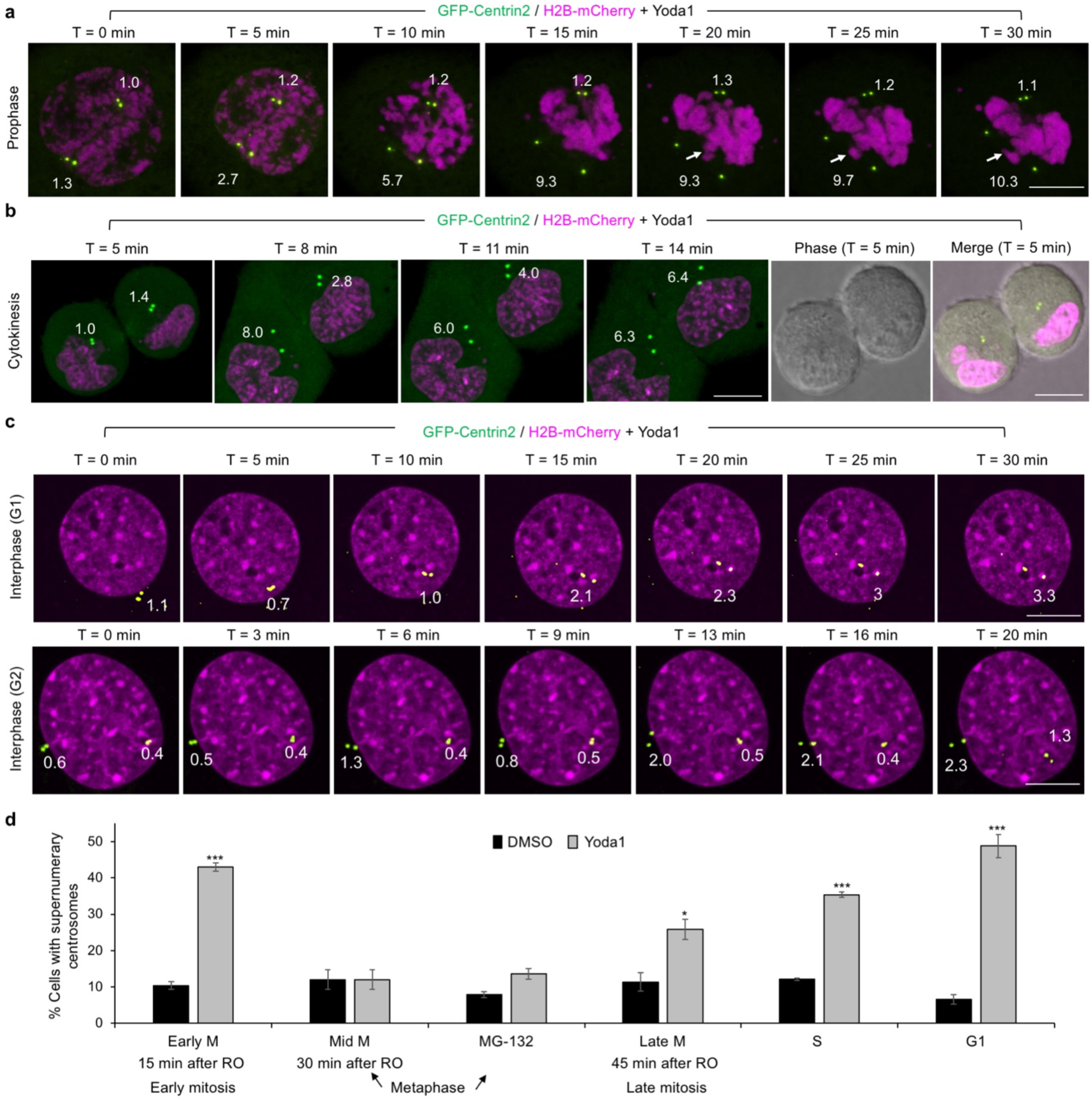
Rapid centriole disengagement following Yoda1 treatment at different cell cycle stages. **a-c,** Live time-lapse images from mitotic or interphase C2C12 cells stably expressing GFP-Centrin2 (green) and H2B-mCherry (magenta) treated with Yoda1 (10 μM). Cells were synchronized to G2/M by RO-3306 (a, b) or interphase by double thymidine block (c). Yoda1 was introduced 15 min and 45 min after RO-3306 washout to capture cells at prophase (a) and cytokinesis (b), respectively, and to unsynchronized interphase C2C12 cells at G1 and G2 (c). Phase images are shown for (b) to emphasis the mitotic stage. Distances (μm) between mother and daughter centriole pairs are marked, and lagging chromatin in (a) is labeled by white arrows. Rapid centriole disengagement was observed at prophase (a), cytokinesis (b) and interphase (c). However, cells at metaphase did not exhibit centriole disengagement upon Yoda1 introduction (Extended Data Fig. 7b). **d,** Quantitative analysis of IF images of C2C12 cells for supernumerary centrosomes following treatment with DMSO (0.1%) or Yoda1 (10 μM) at different stages of the cell cycle, captured largely as in (a-c) and in Extended Data Fig. 7b for metaphase cells. All images are maximum intensity Z projections. All scale bars are 10 μm. Data are represented by mean ± SEM from three independently quantified experiments counting 100-250 cells each. Statistical significance between an experimental group and a control group was assessed by 2-tailed t-test with *** and * for p < 0.0001 and 0.01, respectively.

However, no centriole disengagement was observed for cells at middle mitotic stage (metaphase) (Extended Data Fig. 7b, Supplementary Video 4). We further synchronized cells to metaphase by treatment with MG132, a proteasome inhibitor^43^, and confirmed lack of centriole disengagement at metaphase (Extended Data Fig. 7b, Supplementary Video 8). We also did not observe centriole disengagement events during metaphase in the presence of Dooku1 (Extended Data Fig. 7c, Supplementary Video 9). We speculate that at metaphase, when the chromosomes are already aligned at the spindles, the forces are so strong that Piezo activation will not have an effect at that specific check point. Similar to Yoda1 treatment, we imaged early mitotic cells and interphase cells in the presence of GsMTx4 and observed illegitimate centriole disengagement10-15 minutes after introducing the inhibitor (Extended Data Fig. 7d, Supplementary Video 10-11). Thus, we conclude that both activation and inhibition of Piezo lead to rapid centriole disengagement.

To obtain quantitative data on centriole disengagement and the cell cycle, we also imaged fixed C2C12 cells that were synchronized and treated with Yoda1, and counted cells with supernumerary centrosomes (Fig. 3d, Extended Data Fig. 7e). We detected that the most significant population of cells with supernumerary centrosomes comprises early mitotic cells with 43% of cells at prophase and prometaphase exhibiting supernumerary centrosomes, compared to 10% for DMSO control. Metaphase cells imaged 30 minutes after RO washout or by MG132 synchronization showed no significant supernumerary centrosomes. At late stages of mitosis, 26% of the cells at telophase or undergoing cytokinesis exhibited supernumerary centrosomes compared to 11% for DMSO control. At interphase, 35% of cells synchronized to S phase by double thymidine block exhibited supernumerary centrosomes in comparison to 12% for DMSO control. In addition, 48% of cells synchronized to G1 phase by serum starvation exhibited supernumerary centrosomes in comparison to 7% for DMSO control. Thus, these data indicate that Piezo proteins play an important role in maintaining centrosome integrity not only during interphase but also during mitosis but except metaphase.

### Piezo1 and 2 can mediate Ca^2+^ signaling at centrosomes

During mitosis, extremely high levels of calmodulin (CaM) activation are associated with centrosomes and mitotic spindle poles^44, 45^. Although the source of Ca^2+^ ions at the centrosome is unclear, the localization of Piezo proteins at the centrosome and their ability to transmit Ca^2+^ currents prompted us to consider whether Piezo proteins mediate Ca^2+^ signaling at the centrosome. To monitor local Ca^2+^ levels in individual C2C12 cells, we generated stable cell lines expressing the Ca^2+^-sensitive GCaMP6 reporter targeted to different intracellular locations. GCaMP6 is a highly sensitive Ca^2+^ indicator that represents a fusion between CaM binding peptide (M13), a permuted GFP, and CaM itself^46^, ^47–49^. Upon Ca^2+^ binding, GCaMP6 undergoes a conformational change that increases fluorescence intensity.

One reporter line, carrying a nuclear localization signal (NLS)-tagged GCaMP6 construct, NLS-GCaMP6, exhibited properties especially useful for monitoring centrosomal Ca^2+^ levels. As expected, the NLS-GCaMP6 reporter exhibited increased nuclear fluorescence intensity in response to 1 μM of the Ca^2+^ ionophore ionomycin and to 10 μM Yoda1, and decreased fluorescence in response to the calcium chelator BAPTA-AM (Extended Data Fig. 8a-b). Importantly, after nuclear envelope breakdown in mitosis, strong fluorescence appeared at centrosomes (Fig. 4a) and subsequently around the ends of mitotic spindles (Fig. 4a, Supplementary Video 12). Untagged GCaMP6 reporter also showed strong signals at centrosomes (Extended Data Fig. 8c); so did the Ca^2+^ sensitive dye Fluo4-AM (Extended Data Fig. 8d), supporting the high centrosomal concentration of Ca^2+^. However, the NLS-GCaMP6 line displayed most focused signals at the centrosomes.

**Fig. 4.**
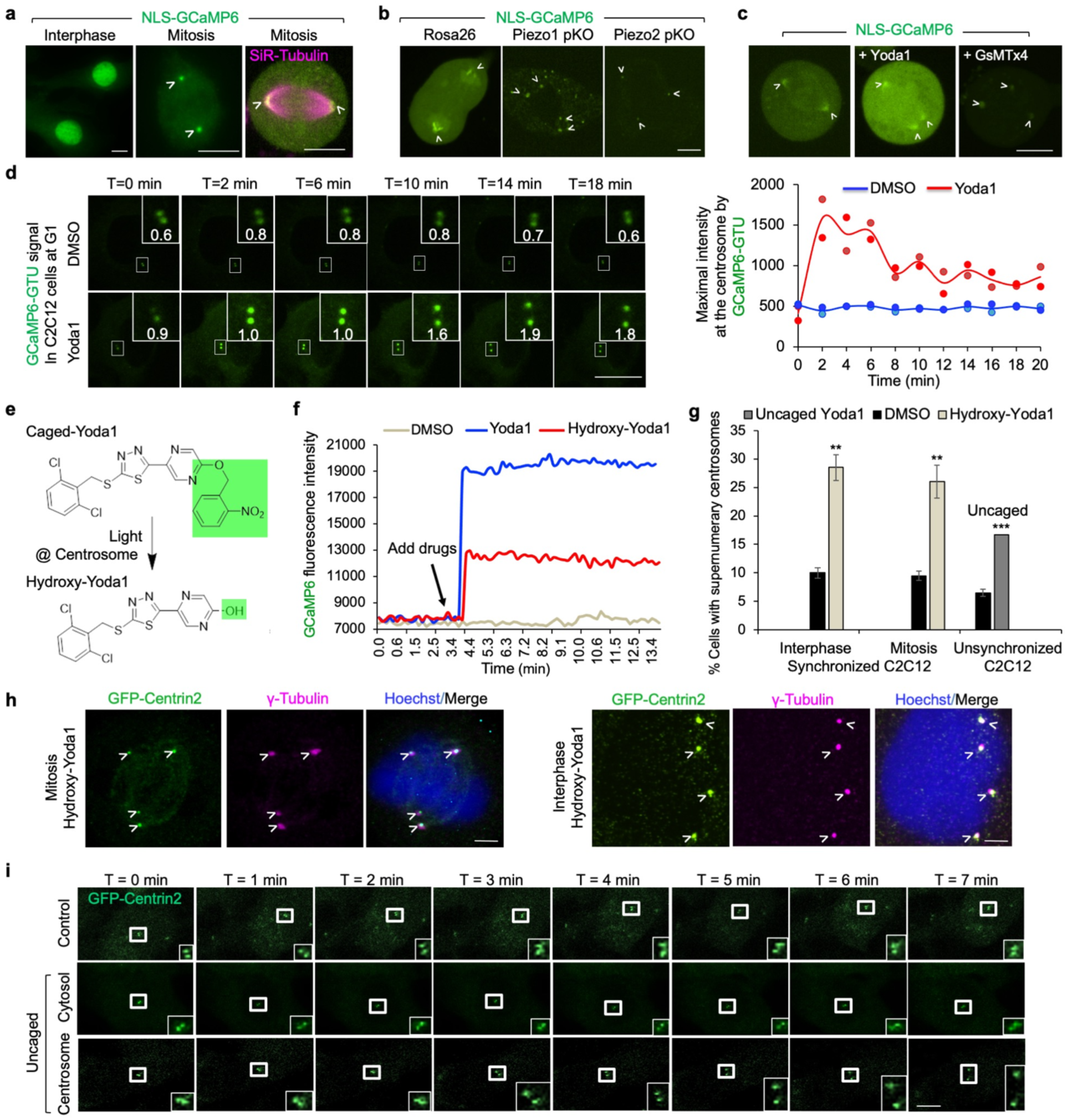
Centriole disengagement induced by specific Yoda1 activation at the centrosome. **a,** Live images of NLS-GCaMP6-C2C12 cells stably expressing an NLS-tagged GCaMP6 Ca^2+^-responsive fluorescence reporter construct. In interphase cells, the NLS-GCaMP6 reporter demonstrates the expected fluorescence in the nucleus *(left)*. Following nuclear envelope breakdown during early mitosis, concentrated fluorescence at the centrosomes was observed (*middle*). In metaphase, the concentrated fluorescence localized to the mitotic spindles, visualized also by SiR-Tubulin-staining of microtubules (magenta) *(right).* **b,** Live images of Piezo1 and 2 pKO NLS-GCaMP6-C2C12 cells compared to an off target Rosa26 pKO control cell. **c,** Live images of NLS-GCaMP6-C2C12 cells, imaged just prior to Yoda1 addition (left), 5 min after Yoda1 (10 μM, middle) or 5 min after GsMTx4 (5 μM, right) addition. In the left and middle panels, the same cell was imaged before and after Yoda1 activation. **d,** Left: live cell imaging of an unsynchronized C2C12 cell (at G1) stably expression GCaMP6-γ-Tubulin (GCaMP6-GTU) after addition of DMSO (top) or Yoda1 (bottom). Upon Yoda1 addition, Ca^2+^ levels at centrosomes increased by ∼ 3 fold, followed by centriole disengagement. Right: average maximal centrosomal Ca^2+^ intensities marked by GCaMP6-GTU (line) as a function of time upon addition of Yoda1 or DMSO. Individual values for the two centrosomes are shown separately as dots. While intensity levels stayed constant with DMSO, they increased with Yoda1 addition and decreased at around the same time when centrioles were separated. **e,** Caged-Yoda1 design that allows photoactivation. The caged group added to Yoda1 is labeled in green. Upon photo uncaging, a hydroxyl group, also marked in green, is formed on the Yoda1 scaffold, generating hydroxy-Yoda1. **f,** NLS-GCaMP6 fluorescence intensities upon addition of hydroxy-Yoda1 (10 μM) as a function of time in comparison to Yoda1 (10 μM) and DMSO. Hydroxy-Yoda1 retained ∼50% of the Yoda1 activity. **g,** Quantitative analysis of percentage of cells with supernumerary centrosomes upon treatment with DMSO or hydroxy-Yoda1 on G2/M synchronized C2C12 cells from IF experiments (h), or upon treatment with caged-Yoda1 followed by light-mediated uncaging on unsynchronized C2C12 cells in comparison with DMSO control from live cell experiments (i). For IF experiments, data are represented by mean ± SEM from three independently quantifications counting 50-200 cells each. The live cell experiments for centriole disengagement by uncaging caged-Yoda1 were performed 36 times with observed 16.7% centriole disengagement. Statistical significance was assessed between an experimental group and the control group by 2-tailed t-test with ***, ** and * for p < 0.0001, 0.001 and 0.01, respectively. **h,** C2C12 cells stably expressing GFP-Centrin2 treated with hydroxy-Yoda1 after release of RO-3306, imaged by IF after 30 min for Centrin2 (GFP fluorescence), γ-Tubulin (magenta) and DNA (Hoechst dye, blue). Supernumerary centrosomes and centriole disengagement are shown in mitotic and interphase cells. **i,** An example of centriole disengagement upon uncaging caged-Yoda1 at the centrosome (bottom row) in comparison with controls. C2C12 cells stably expressing GFP-Centrin2 and treated with 10 μM caged-Yoda1 were imaged after application of photolytic light at centrosomes (bottom row), away from centrosomes (middle row) or no application (top row) at T = 0 min. Selective snapshots are shown with 1 min intervals. The cells at the top and bottom rows are from the same movie (Supplementary Video 15), but only the light-treated centrosome underwent prompt centriole disengagement.

When imaged on day 1 after Piezo pKO selection, Piezo1 and 2 C2C12 pKO cells exhibited reductions in maximal NLS-GCaMP6 fluorescence intensity at centrosomes to 44% and 34%, respectively, of the average level in Rosa26 control cells (Fig. 4b, Extended Data Fig. 8e). The difference in fluorescence was not due to changes in GCaMP6 protein levels at the centrosome, as anti-GFP IF showed similar levels of GCaMP6 protein localization at centrosomes in Rosa26 and Piezo pKO C2C12 cells (Extended Data Fig. 8f). Treatment with GsMTx4 reduced GCaMP6 fluorescence to 60% of the level before treatment (Fig. 4c, Extended Data Fig. 8g). We also imaged Ca^2+^-induced fluorescence intensity at centrosomes in mitotic GCaMP6 C2C12 cells 5 minutes after Yoda1 addition. Upon introducing Yoda1, maximum fluorescence intensity at centrosomes and spindle poles increased by 2.2-fold compared to the level before stimulation (Fig. 4c, Extended Data Fig. 8g). Of note, GCaMP6 signal in the cytosol also increased proportionally, likely due to Piezo distribution in other intracellular compartments (Extended Data Fig. 8g).

In order to be able to track calcium local concentration specifically at the centrosome upon Piezo1 alterations, we designed and prepared C2C12 stable line expressing GCaMP6 fused to γ-Tubulin (GCaMP6-GTU). We performed live cell imaging of unsynchronized C2C12 cells at G1 in the presence of Yoda1 or DMSO (Fig. 4d. Supplementary Video 13, 14). We noticed that while calcium levels stayed constant in the presence of DMSO, they increased by ∼3 fold within a few minutes upon Yoda1 addition and decreased at ∼10 min when centrioles were also separated. Because of the GTU localization, this GCaMP6 reporter did not show much increase in the cytosolic signal. Collectively, these observations support a model whereby Piezo1 and 2 balance local Ca^2+^ levels at the centrosomes to prevent inappropriate centriole disengagement and supernumerary centrosome formation.

### Specific Yoda1 uncaging at the centrosome induced illegitimate centriole disengagement

The ability of cell permeable Yoda1, but not cell impermeable Jedi1, Jedi2 or ionomycin to induce supernumerary centrosomes suggests that centrosomal Piezo1 mediates this effect, potentially by modulating local Ca^2+^ concentration at the centrosome. This hypothesis is supported by the sufficiency of Ca^2+^ to activate centriole disengagement in a cell free system^50^. To test the role of centrosomal Piezo1 directly, we designed and synthesized a modified “caged”-Yoda1 that is normally inactive but can be uncaged or activated specifically at centrosomes by applying 405 nm photolytic light to GFP-Centrin2 marked centrosomes via a confocal microscope to release hydroxy-Yoda1 (Fig. 4e, Extended Data Fig. 9a). The activity of hydroxy-Yoda1 was first tested in C2C12 cells stably expressing NLS-GCaMP6. The levels of GCaMP6 fluorescence intensity, reflecting local Ca^2+^ concentrations, were measured after introducing Yoda1, hydroxy-Yoda1 or DMSO as control. Hydroxy-Yoda1 showed ∼50% of the activity of Yoda1 in activating Ca^2+^- induced fluorescence (Fig. 4f). We then verified by IF that like Yoda1 although with reduced activity, hydroxy-Yoda1 caused centriole disengagement in 29% and 26% interphase and mitotic cells, respectively (Fig. 4g-h).

Next, we imaged unsynchronized C2C12 cells stably expressing GFP-Centrin2 to observe whether illegitimate centriole disengagement occurred upon Yoda1 uncaging. Upon photolytic uncaging of Yoda1 at centrosomes, we observed that centrioles separated to > 5 μm in distance within 3 minutes by live cell imaging and stayed apart throughout the imaging period (Fig. 4i, Supplementary Video 15, 16). By contrast, control cells in the same field containing caged-Yoda1 that was not uncaged by photoactivation or control cells uncaged not at the centrosome showed normal centrosome behavior (Fig. 4i, Supplementary Video 15, 17). To obtain statistics on uncaging-induced centriole disengagement, we performed the experiment 36 times, and captured rapid centriole disengagement 6 times, resulting in 16.7% cells with supernumerary centrosomes, significantly different from the 6.5% in unsynchronized DMSO control cells (Fig. 4g).

Of note, caged-Yoda1 activation at the centrosome did not cause cell surface protrusions, suggesting lack of plasma membrane Piezo1 activation (Supplementary Video 16). Collectively, these Yoda1 uncaging experiments directly support the idea that Yoda1-induced centriole disengagement is mediated by the pericentrosomal pool of Piezo1 and that Piezo proteins may sense microtubule forces at centrosomes in order to maintain their integrity.

### Centrosomal endosomes recruit Piezo by Snx5

Piezo1 and 2 are *bona fide* transmembrane proteins; however, the centrosome is a conventional membraneless organelle. To identify the membrane compartment in the centrosomal region in which Piezo proteins presumably reside, we conducted a yeast two-hybrid (Y2H) screen to interrogate a prey library from adult and fetal human skeletal muscle. Because of its striking evolutionary conservation and the fact that most missense mutations in *PIEZO1* and *2* GOF disease states cluster in the 70 aa Piezo C-terminal domain (CTD)^16^, we used the Piezo2 CTD as bait. In this initial Y2H screen, we identified the sorting nexin, Snx5, as a top candidate (Extended Data Fig. 9b). Sorting nexins, including Snx5, are responsible for trafficking endosomes to specific intracellular destinations^51–56^. They often bind to C-termini of transmembrane endosome cargo proteins that protrude from endosome membranes into the cytoplasm^57^. Consistent with an Snx5 interaction, cryo-EM structures of Piezo1 and Piezo2^58–62^ predict a cytosolic orientation for the Piezo CTD. Importantly, recycling endosomes trafficked by Snx5 have been shown by TEM to reside in very close proximity to the centrosome via microtubule-dependent retrograde transport^51, 53^

The interaction between the Piezo2 CTD and Snx5 was validated using a different Y2H system from that employed in the original screening (Fig. 5a). Immunoprecipitation followed by Western blot using epitope-tagged Piezo2 and Snx5 constructs transfected into HEK293T cells confirmed the interaction (Fig. 5b). In addition, IF imaging of Snx5 in C2C12 and Neuro-2A cells demonstrated co-localization with both Piezo2 and γ-Tubulin (Fig. 5c). We next isolated centrosomes from C2C12 cells, fractionated them using a sucrose gradient, and assayed the resulting fractions by Western blots for Piezo1 and Piezo2, for γ-Tubulin and Pericentrin as centrosome markers, and for Snx5 as a marker specific to the endosomal class of interest (Fig. 5d). Piezo1 and 2 fractions highly overlapped with Snx5 fractions, while Pericentrin and γ-Tubulin partially overlapped with Piezo1, Piezo2 and Snx5. Of note, Pericentrin has been identified from an anti-Piezo2 immunoprecipitate as a Piezo2-interacting protein^63^.

**Fig. 5.**
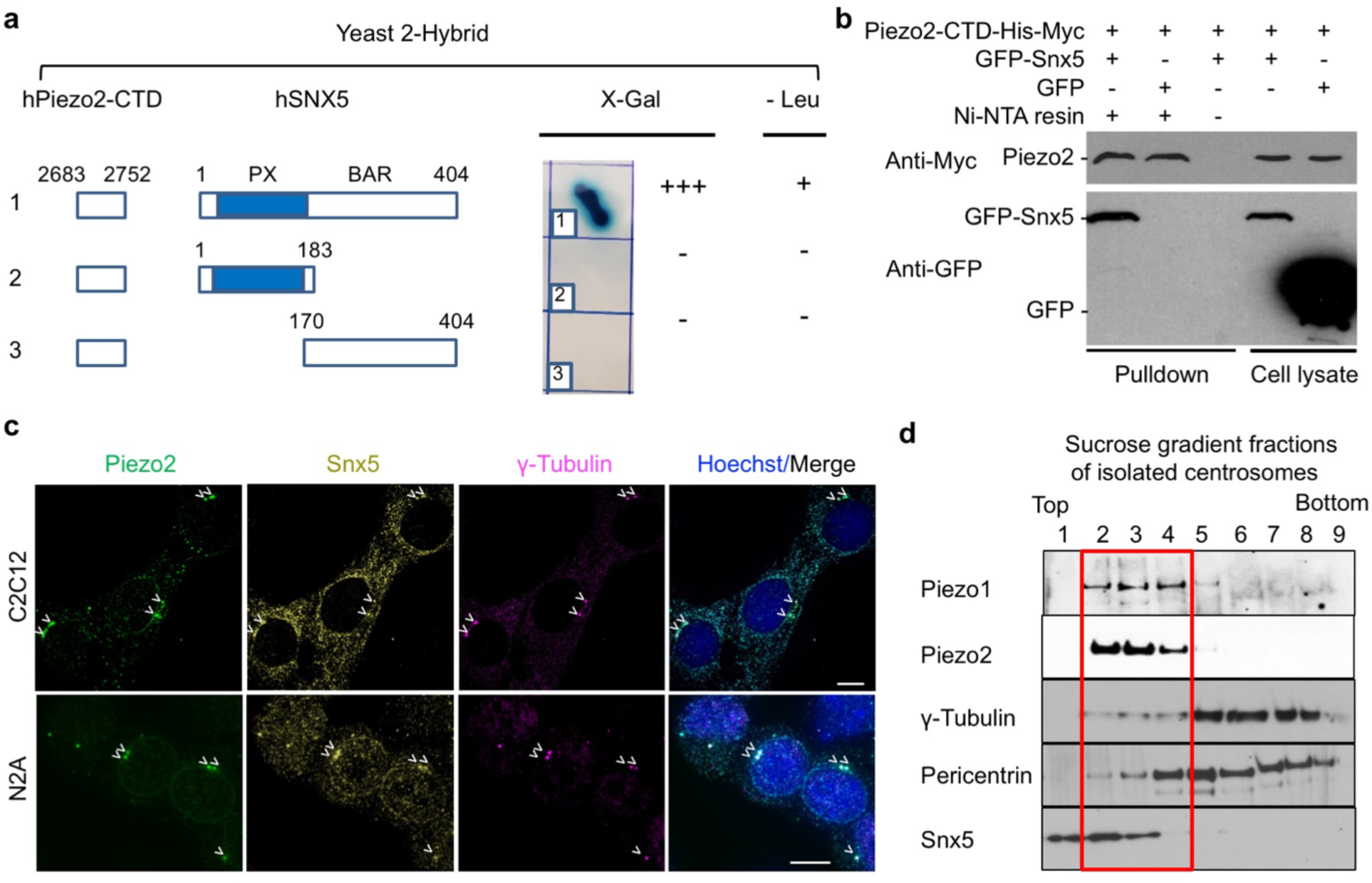
Piezo endosomal localization. **a,** Validation of the Snx5 interaction with the Piezo2 C-terminal domain (CTD, last 70 aa) using an alternative yeast two-hybrid assay. The Piezo2 CTD was fused to the LexA DNA binding domain and full-length Snx5 was fused to a B42 activation domain. Positivity is indicated by yeast turning dark blue in the presence of the LacZ reporter X-gal (+++) and by yeast growth in the absence of leucine. Negativity is indicated by yeast staying white (-) and no growth in the absence of leucine. Phox homology (PX) domain for lipid binding and Bin/Amphiphysin/Rvs (BAR) domain for membrane-curvature sensing are shown. **b,** Western blot analysis of Ni-NTA pulldown of human Snx5 by human Piezo2 CTD. Piezo2-CTD-His-Myc was co-expressed with GFP-Snx5 or GFP control in HEK293T cells. The His-tagged protein was precipitated using Ni-NTA beads (lane 1&2) or as a control with NTA beads without Ni (lane 3). Lanes 4 and 5 are cell lysates. Precipitated Piezo2-His-Myc was detected with anti-Myc antibody and GFP-hSnx5 was detected with anti-GFP antibody. **c,** IF images of C2C12 cells (top panel) and Neuro-2A cells (bottom panel) stained for Piezo2 (green), Snx5 (olive), ɣ-Tubulin (magenta) and DNA (blue). Co-localization among Piezo2, Snx5 and ɣ-Tubulin is shown by arrowheads and all scale bars are 10 μm. **d,** Western blot analysis of fractionation of isolated centrosomes. Centrosomes isolated from C2C12 cells were fractionated on a sucrose gradient, and blotted with the following antibodies: anti-Piezo1, anti-Piezo2, anti-γ-Tubulin, anti-Pericentrin and anti-Snx5.

We further used microtubule disrupters to investigate Piezo localization. Because nocodazole, a frequently used drug that interferes with microtubule polymerization, is toxic to C2C12 cells, we tested the effect of Parthenolide, which interferes with microtubule structures via multiple mechanisms^64^. Treatment with Parthenolide showed reduced or no observed Piezo1/2 localization at centrosomes (Extended Data Fig. 10a, b). 3h cold treatment for destabilizing microtubules resulted in reduced Piezo1 localization at centrosomes however, had no apparent effect on the Piezo2 localization (Extended Data Fig. 10a, b). By contrast, treatment by Taxol, a microtubule stabilizer, had no apparent effect on the Piezo1/2 localization (Extended Data Fig. 10a, b). These data collectively support that Piezo1 and 2 on endosomes are transported to centrosomes by microtubule retrograde transport.

### PLK1 inhibition abolished Yoda1-induced centriole disengagement

In order to obtain a deeper mechanistic understanding on Piezo activity modulation-induced centriole disengagement, we tested the role of Polo-like-kinase 1 (PLK1), a serin/threonine kinase that has a recognized role in coordinating procentriole maturation and centriole duplication^50, 65–68^. We first found by Western blots that in C2C12 cells, PLK1 expression was high in G2, low in S and M and lowest in G1 (Extended Data Fig. 10c); Piezo2 appeared to follow the PLK1 expression pattern by cell cycle stages while Piezo1 expression was mostly constant throughout cell cycle with a slight enrichment in G1 (Extended Data Fig. 10c). Of note, these experiments were performed by synchronization to G1, S, G2 and M using, respectively, 24 h serum starvation, double thymidine block, 12 h RO treatment, and plate shake off at 30 min after RO washout.

We titrated different concentrations of BI-6727 (Volasertib, 1 nM to 100 nM), a highly selective and potent PLK1 inhibitor^69^, in C2C12 cells, performed cell cycle analysis based on propidium iodide (PI) staining and determined the optimal working concentration in C2C12 cells as 100 nM (Extended Data Fig. 10d). Incubation of C2C12 cells with 100 nM BI-6727 for 18 h followed by 30 min treatment with DMSO, Yoda1 or GsTMx4 showed that BI-6727 prevented the appearance of supernumerary centrosomes shown by IF and quantification (Fig. 6a, b). We also used IF to determine the effect of Piezo1 activation by Yoda1 on PLK1 distribution and found that PLK1 partially localized on the centrosome either before or after Yoda1 treatment (Fig. 6c). These data suggested that Piezo1 and 2 impact centrosomes by acting upstream of PLK1 to regulate PLK1 activity.

**Fig. 6.**
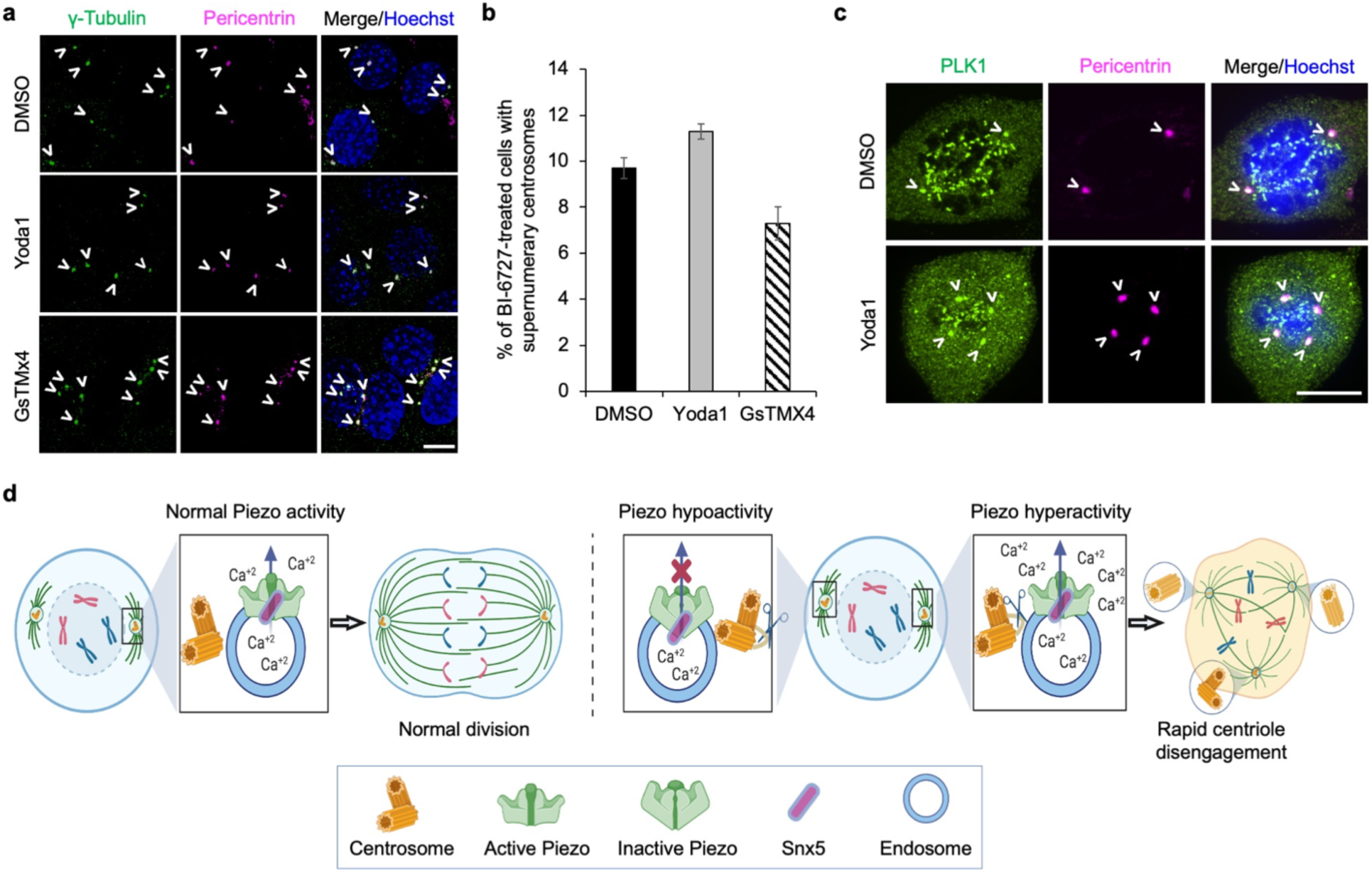
PLK1 inhibition prevents supernumerary centrosomes. **a**, IF of C2C12 cells stained with γ-Tubulin (green), Pericentrin (magenta) and DNA (blue) after treatment with the PLK1 inhibitor BI-6727 for 18 h and with Yoda1, GsTMx4 or DMSO (control) for 30 min. **b**, Quantification of cell populations with supernumerary centrosomes for IF images in (a). **c,** IF images of mitotic C2C12 cells stained for PLK1 (green), Pericentrin (magenta) and DNA (blue), fixed 30 minutes after addition of DMSO (control) or Yoda1. No apparent changes in PLK1 localization at centrosomes were observed. Scale bars: 10 μm and centrosomes are marked with white arrowheads. **d,** Proposed model for how Piezo-mediated Ca^2+^ signaling regulates centriole disengagement. Piezo-containing endosomes or other vesicles near centrosomes may respond to mechanical forces transmitted by microtubules that deform the vesicular membrane, activating Piezo to release internally stored Ca^2+^ and activating Ca^2+^-sensitive centrosomal proteins that maintain centrosome integrity. Normal, hyperactivated, and hypoactivated Piezo states are shown. Either Ca^2+^ excess or deficit leads to rapid centriole disengagement.

Interestingly, a similar centriole phenotype dependent on PLK1 activity was also observed after a prolonged RO treatment of 48 h^67^. Because PLK1 activity during early mitosis is actually required for subsequent disengagement of centrioles at late mitosis ^67, 70^, mitotic delay- and Yoda1- induced phenotypes, one slow and one fast respectively, may be related to the same or similar downstream pathways in the “normal” centrosome cycle. Furthermore, it was shown that PLK1 activity during late mitosis promotes centriole disorientation during G1 in order to allow reduplication at S phase^66, 67^. This may provide an explanation to Yoda1-induced centriole disengagement at G1, despite low expression of PLK1 at G1. However, centriole disengagement in G1 may also be a very different process as centrioles are already disorientated due to the loss of their orthogonal confirmation^65^.

## Discussion

The present study demonstrates that in addition to its known role in sensing forces that impinge on the cell membrane, Piezo signal transduction also operates intracellularly in the context of centrosome function (Fig. 6d). We provide evidence that Piezo proteins exhibit pericentrosomal localization, likely through an association with endosomes that dynamically localize to the centrosome^52, 53, 56, 71^ and that both Piezo hyperactivation and inhibition can induce supernumerary centrosomes. This intriguing result is potentially consistent with the observation of similar arthrogryposis phenotypes in humans carrying either *PIEZO2* gain- and loss-of-function mutations^17–21^. This fundamental cellular function of Piezo proteins is also supported by the embryonic and perinatal lethality, respectively, in *Piezo1*^-/-^ and *Piezo2*^-/-^ mice^8, 23^. Previous studies have implicated force generation by microtubules as essential for maintaining centrosomal integrity^41^, but the proteins that mediate this mechanotransduction have remained unknown. We propose that Piezo proteins are key regulators of this evolutionarily conserved process, which is important not only for normal cell cycle, but also for aberrations such as cancer which is frequently associated with supernumerary centrosomes. Is it possible that balanced cytoskeletal forces with optimal Piezo-induced local Ca^2+^ signals may act upon the pericentriolar matrix and the centrosome to regulate centrosome integrity during the cell cycle^72^? In this regard, many centrosomal proteins that regulate the centrosome pathway are known to be Ca^2+^-sensitive, including Centrins, Pericentrin, Separase and CaM^73–77^. A full understanding of how Piezo mechanotransduction and Ca^2+^ signaling act at the centrosome, or intracellularly in general, will undoubtedly provide fresh insights into Piezo-mediated human diseases.

## Acknowledgements

We thank Iain Cheeseman and Ardem Patapoutian for discussions, Paula Montero Llopis and Microscopy Resources on the North Quad (MicRoN) core at Harvard Medical School for microscope use and training, Maria Ericsson, Margaret Coughlin and Anja Nordstrom from the Harvard Medical School EM Core for cell sectioning for TEM and SEM and assistance with EM imaging, Margaret Muscato, Ken Anderson and Steve Coleman from Biovis for their assistance with iSIM imaging, Emily Cronin-Furman, Erin Powrie and Kathy Lindsley from Olympus for their assistance with STORM imaging, and Grigoriy Losyev at the flow cytometry core at Brigham and Women’s Hospital for assistance with cell sorting. This work was supported by NIH grants U01 DE024443 and U01 HG007690 (R.L.M.) and by a fellowship from the Cancer Research Institute (L.D.).

## Author contributions

L.D., H.W., and R.L.M. conceived the project idea and designed the study. L.D. designed and cloned constructs. L.D prepared stable cells. L.D. and K.A.K. designed and cloned gRNAs for CRISPR-Cas9 KO. L.D. and L.M. performed CRISPR-Cas9 KO experiments. L.M. isolated and cultured primary myoblasts. L.M. performed Immunofluorescence and confocal imaging for primary myoblasts. L.D and L.M performed cell cycle analysis for primary myoblasts and C2C12 cells. L.D. performed Immunofluorescence and confocal imaging for experiments in cell lines. Q.X., L.M. and L.D. prepared mouse sections and performed confocal imaging for antibody validation in tissues. Y.Z. performed shRNA experiments. J.V.S. assisted with Piezo1-GFP imaging and provided suggestions. L.D. performed electron microscopy for cell sections. L.D. performed calcium imaging and live cell imaging. L.D. designed and performed caged-Yoda1 experiments. X.F performed Y2H validation and Pulldown for Piezo2 and Snx5. L.D performed organelle fractionation and western blots. L.D. and L.M. performed image analysis and statistics. L.D., H.W. and R.L.M wrote the manuscript with input from all authors.

## Methods

### Constructs and cell culture

The different cell lines used in this study (Neuro-2A, C2C12, IMCD3 and HEK293T) were cultured in Dulbecco’s modified Eagle’s medium (DMEM) supplemented with 10% fetal bovine serum (FBS). Full-length mouse Piezo1 was fused with an eGFP-FLAG C-terminal tag and cloned into the pCDNA3.1 vector containing a neomycin resistance cassette to generate the Piezo1-GFP construct. The eGFP fused Centrin2 (GFP-Centrin2, Addgene) and Lenti-H2B-mCherry (GeneCopoeia) plasmids were purchased.

For preparing cell lines stably expressing Piezo1-GFP, GFP-Centrin2, GCaMP6s-GTU or NLS-GCaMP6s, C2C12 cells were transfected with the appropriate construct using lipofectamine 3000 (Thermo Fisher Scientific). Forty-eight h after transfection, the culture medium was replaced by selection medium containing DMEM, 10% FBS and 500 μg/ml G418 (Thermo Fisher). Cells were kept under selection for 2 weeks, whereas all untransfected cells cultured in parallel under the same selection died after 6 days. For generating a C2C12 cell line stably expressing both GFP- Centrin2 and H2B-mCherry, C2C12 GFP-Centrin2 cells were infected with Lenti-H2B-mCherry and sorted after 2 days of selection in 2 μg/ml puromycin. Surviving positive cells were sorted using a BD *FACS*Aria^TM^ Sorter at the Brigham and Women’s Hospital Genetics Division Flow Cytometry Core. Prior to live cell or fixed cell imaging, positive cells were re-sorted to further enrich populations of interest.

### Mice

*Piezo2*^-/-^ mice generated by breeding *Piezo2^fl/fl^* and *EIIaCre* mice^14^ were purchased from The Jackson Laboratory (stock 027720).

### Primary myoblast isolation and cell culture

Isolation of primary mouse myoblasts was performed as previously described^30^ with minor adaptations. Briefly, forelimbs and hindlimbs were removed from newborn (P0) mice and the bones and fibrotic tissue were dissected away. The remaining muscle mass was weighed and then briefly immersed in 70% ethanol and washed with fresh Dulbecco’s phosphate buffered saline (DPBS). For each 20 mg of tissue, 2 mL of a mixed solution containing 1.5 U/ml collagenase type 2 (Worthington LS004174), 2.4 U/ml dispase II (Sigma-Aldrich D4693), and 2.5 mM CaCl_2_ in serum-free DMEM (Sigma-Aldrich) was used to dissociate cells from muscle tissue that had been minced into a slurry using sterile razor blades. The slurry was incubated at 37 °C for 30 min, triturated every 10 min with a pipette, and then passed through a 70 μm nylon mesh. The filtrate was spun at 500 x g for 5 min to pellet the cells that were then resuspended in Ham’s F-10 growth media (Thermo Fisher), 20% FBS, 1% penicillin/streptomycin, 1% α-glutamine and 2.5 ng/ml of βFGF (PeproTech) and plated on dishes coated using 10 μg/ml laminin (Sigma-Aldrich). To increase myoblast population purity, a 30-min pre-plating was performed at the first passage. After two passages, the culture was enriched for myoblasts based on MyoD staining. The media was then switched for subsequent passages to growth media (high-glucose DMEM complemented with 20% FBS, 10% horse serum, 1% penicillin/streptomycin, 1% α-glutamine).

### CRISPR-Cas9 polyclonal knockout

The following two Piezo sgRNAs and a control sgRNA were cloned into the lentiCRISPRv2 vector (Addgene) containing Cas9, tested in C2C12 cells and validated by Western blotting.

Piezo1 exon 8 sgRNA: 5’-GTGCTTAGTGTTGAGTACCA-3’
Piezo2 exon 38 sgRNA: 5’-GGACTGCCTTGAGATCAGCA-3’
Rosa26 Off Target control sgRNA: 5’-GACTCCAGTCTTTCTAGAAGA-3’

For virus production, we used lentiviral packaging plasmid pCMV Delta R8.2 (Addgene) and the envelope plasmid pCMV-VSV-G (Addgene). For each sgRNA, HEK293T cells plated on 10 cm plates were cultured in DMEM supplemented with 10% FBS and transfected using lipofectamine 3000 with 5 μg lentiCRISPRv2 plasmid, 3.3 μg packaging plasmid and 1.7 μg envelope plasmid. Forty-eight h post-transfection, viruses were collected and used on the same day for infection of C2C12 cells. C2C12 cells were incubated for 3 h with 8 μg/ml polybrene prior to introduction of the viruses at a ratio of 1:4. One day post infection, cells were selected with puromycin at a final concentration of 2 μg/ml for 2 days.

### Piezo lentiviral shRNA knockdown in C2C12 myoblasts

shRNA MISSION TRC-Mm 2.0 (Mouse) RNA interference lentiviruses targeting mouse Piezo2 were obtained through the Broad Institute. The shRNA targeting sequences are show below.

Control Off Target: 5’-AGAATCGTCGTATGCAGTGAA-3’ (TRCN0000072250)
Piezo1 sh #1 (Ex14): 5’-CACCGGCATCTACGTCAAATA-3’
Piezo1 sh #2 (Ex18): 5’-TCGGCGCTTGCTAGAACTTCA-3’
Piezo2 sh #1 (Ex47): 5’-GAGGACATTTACGCGCACATT-3’ (TRCN0000125186)
Piezo2 sh #2 (Ex46): 5’-CCTCACAAAGAGCTACAATTA -3’ (TRCN0000125185)
Piezo2 sh #3 (Ex15): 5’-TCATGAAGGTGCTGGGTAA-3’

For lentivirus infection, C2C12 myoblasts were plated in 6 well plates at a density of 2 × 10^5^ cells/dish 24 h prior to infection with viral media. Before transduction, cell medium was replaced with fresh medium containing polybrene (8 μg/ml). Lentiviruses were then added to the cell plates were spun at 1000 x g for 30 min. After overnight incubation (24 hr), lentiviruses were removed, cells were washed with PBS, and medium containing puromycin (2 μg/ml) was added for drug selection for 48 h.

### Quantitative real-time PCR (qRT-PCR)

Total RNA from shRNA transduced C2C12 cells was extracted using RNAqueous®-Micro Total RNA Isolation Kit (ThermoFisher). 1 μg total RNA was used to generate one strand cDNA using the qScript™ cDNA Synthesis Kit (Quanta Biosciences). Real time Taqman PCR assays for mouse Piezo2 were purchased from Applied Biosystems with a FAM reporter dye and a non-fluorescent quencher. Universal TaqMan PCR master mix (20X) with AmpErase UNG (Thermo Fisher Scientific) was used. The reaction was run on a LightCycler (Bio-Rad) using 1 μl of the cDNA in a 20 μl reaction according to the manufacturer’s instructions in triplicate. Calibrations and normalizations were done using the 2^-ΔΔCT^ methods. The reference gene used was GAPDH.

### Immunofluorescence (IF)

Cell lines and primary myoblasts were plated on LabTek II 4-well chamber slides, LabTek 35 mm bottom glass dishes or CELLview 4-compartment dishes (Greiner Bio-One), fixed in 100% methanol for 5 minutes at -20 °C or 4% paraformaldehyde (PFA) for 10 min at room temperature and permeabilized with 0.1% Triton X-100 for 10 minutes. Cells were incubated in PBS with Tween 20 (PBST) containing 10% normal goat serum for 1 h, which minimized non-specific binding. After three washes with PBST, cells were incubated overnight at 4 °C with primary antibody. The following antibodies were used for Piezo detection: rabbit polyclonal anti-Piezo1 antibody (1:200 Novus Biologicals NBP1-78446) and rabbit polyclonal anti-Piezo2 antibody (1:200 Novus Biologicals NBP1-78624). For other markers the following antibodies were used: rabbit polyclonal Pericentrin (1:500 Abcam ab4448), rabbit polyclonal Cep164 (1:200 Abcam ab221447), mouse monoclonal anti PLK1 (1:500, Abcam ab17056), mouse monoclonal anti-γ-Tubulin (1:500 Abcam ab11316), or rabbit polyclonal anti-α-Tubulin (1:1000 Abcam ab18251). After incubation, cells were washed and incubated with Alexa Fluor 647-conjugated anti-rabbit IgG (1:500) and Alexa Fluor 488-conjugated anti-mouse IgG (1:500) or Fluor 568-conjugated anti-mouse IgG (1:500) for one h at room temperature, washed with PBS (3 x 10 min) and then stained with Hoechst (1:1000). For negative control we used rabbit IgG (Dako) at the same concentration for Piezo primary antibodies. Cells were imaged using a Nikon Ti inverted microscope fitted with a Photometrics CoolSNAP HQ2 Peltier cooled CCD camera and Andor Zyla 4.2 sCMOS camera equipped with Plan Apo 100x/1.4 Ph3, Plan Apo 60x/1.3 DIC and Plan Apo 40x/1.3 DIC objectives. Lumencor SpectraX LED illuminator was used. Chroma 49000 (DAPI), Chroma 29002 (green), Chroma 49008 (red) and Chroma 49011 (far red) filter cubes were used. Image analysis was performed in Fiji^78^.

### Immunofluorescence (IF) of mouse sections

All tissues were briefly fixed in 4% PFA in PBS and then incubated in 15% sucrose in PBS for 4 h followed by incubation with 30% sucrose in PBS overnight at 4 °C. Fixed tissues were cryo-embedded in OCT compound (Sakura) and cryo-sectioned for immunofluorescence. Muscles and DRG were sectioned from mouse embryos and newborn mouse pups at the indicated stages. Lungs were removed from newborn mouse pups. For IF analysis, mice sections were incubated with primary antibodies diluted in blocking solution (5% goat serum in PBST) for 18 h at 4 °C and with secondary antibodies conjugated to Alexa fluorophores (488, 647) for 2 h at RT. Specimens were stained with Hoechst and imaged with a Zeiss 760 or Nikon Spinning disk confocal microscope.

### Cell cycle analysis

At 70% confluence, C2C12 cells and mouse primary myoblasts were trypsinized, washed with PBS and fixed in ice cold 70% ethanol for 30 min at 4 °C. Cells were then washed with PBS and stained with propidium iodide (PI) (Sigma-Aldrich) and analyzed on a Cytek DxP 10 flow cytometer using the Brigham and Women’s Hospital Division of Genetics Flow Cytometry Core. Cell cycle distribution was analyzed using Flow Jo software.

### Calcium imaging

GCaMP6s was cloned into the pcDNA3 vector with the CMV promoter (a gift from Dr. David Clapham, HHMI), as a fusion at its N-terminus to a nuclear localization sequence (PKKKRKV). GCaMP6s-GTU construct was made based on mCherry-**γ**-Tubulin-17 backbone (Addgene #55050). GCaMP6s constructs were stably expressed in C2C12 cells to perform Ca^2+^ imaging. Validation of the Ca^2+^ reporter function was performed with 1 μM ionomycin (Sigma) to increase calcium flux, and a small-molecule calcium chelator, 1,2- bis(2-aminophenoxy)ethane-N,N,N ′, N ′ -tetraacetic acid tetrakis (acetoxymethyl ester) (BAPTA-AM) (Sigma) was used at 10 μM to decrease calcium level. For Ca^2+^ imaging with the Fluo4-AM dye (Thermo Fisher Scientific), cells were loaded with 9 μM Fluo4-AM solubilized in DMSO and supplemented with 1% Pluronic F-127 by incubating at 37 °C for 45 min. After loading, cells were incubated with fresh medium for 20 min before starting to image in order to allow for complete de-esterification of the AM ester.

### Live cell imaging

Live cell imaging of C2C12 cells stably expressing Piezo1-GFP was performed on a Nikon confocal microscope at 20x magnification with 1.5 zoom for up to 24 h at 5 min intervals. For live cell imaging of C2C12 cells stably expressing GFP-Centrin2 and H2B-mCherry, or C2C12 cells stably expressing NLS-GCaMP6 during mitosis, cells were plated on LabTek 35 mm bottom glass or CELLview 4-compartment dishes and imaged while cultured in Hank’s balanced salt solution (HBSS) supplemented with 10% FBS and 1% penicillin/streptomycin at 37 °C with 5% CO_2_ using a spinning disk confocal Nikon microscope equipped with temperature controller and CO_2_ incubator, using Plan Apo 60x/1.3 oil objective. Analyses of live cell imaging data was performed in Fiji^78^.

### Live cell imaging for caged-Yoda1 upon uncaging and hydroxy-Yoda1 activity

The caged-Yoda1 and hydroxy-Yoda1 were synthesized by WuXi AppTec with 98% purity and characterized by nuclear magnetic resonance and liquid chromatography-mass spectrometry. Live cell imaging of C2C12 cells stably expressing GFP-Centrin2 C2C12 was performed on a Zeiss confocal microscope equipped with temperature controller and CO_2_ incubator using a 63x objective. Once cells were treated with 10 μM caged-Yoda1, we uncaged Yoda1 by applying photolytic light (405 nm laser, 100 μsec, 20 iterations) of ∼1.5 micron in diameter at the centrosomes, followed by imaging cells for 20 min at 8 sec intervals, or 10 min at 20 sec intervals. Hydroxy-Yoda1 activity was measured at 10 μM in comparison to 10 μM Yoda1 and a DMSO control by plotting the fluorescence intensity (excitation at 488 nm and emission at 520 nm) of NLS-GCaMP6 cells using a Neo BioTek plate reader.

### Super-resolution microscopy

We performed instant structured illumination microscopy (iSIM) for unsynchronized C2C12 cells stained for Piezo2 (Alexa 647 conjugated) and γ-Tubulin (Alexa 568 conjugated) using Biovision VT-iSIM Super Resolution System on a Leica inverted DMI8 microscope equipped with a Quest camara (Hamamatsu), a Ludl X, Y motorized stage with a Ludl piezo and a 100x objective. We also performed stochastic optical reconstruction microscopy (STORM) for unsynchronized C2C12 cells using an Olympus Abbelight SAFe 360 system on an IX83 Olympus inverted microscope equipped with a Hamamatsu Fusion camera and a 100x objective. Image deconvolution was performed with the Abbelight Neo SAFe software.

### Western blot analysis

Western blot analysis was performed for WT C2C12 cells, Piezo1 pKO, Piezo2 pKO and Rosa26 off target C2C12 cells. Samples were loaded on a NUPAGE 3-8% Tris acetate gel (Thermo Fisher Scientific) or 4-15% mini PROTEAN Tris-Glycine gel (Bio-Rad), and the gel was transferred to a nitrocellulose membrane by using wet transfer for 3 h at 90 mV or transfer for 10 min at 20 mV with Invitrogen iblot 2 gel transfer device. After transfer, the membrane was blotted with 5% non-fat milk in Tris-buffered saline with Tween 20 (TBST) for 1 h, followed by overnight incubation with either Piezo1 rabbit polyclonal antibody (1:1000, Proteintech 15939 1-AP), Piezo2 rabbit polyclonal antibody (1:1000, Novus Biologicals NBP1-89892), PLK1 mouse monoclonal (1:1000 Abcam ab17056), GAPDH mouse monoclonal antibody (1:5000, Proteintech 60004-1) at 4 °C. After incubation with primary antibody, the membrane was washed 3 times with TBST and incubated for 1 h with Peroxidase AffiniPure goat anti-rabbit IgG or goat anti-mouse IgG (Jackson Laboratories). The membrane was developed on film or Bio-Rad gel imager with enhanced chemiluminescence (ECL) detection (Thermo Fisher Scientific). Densitometric analysis of protein bands was performed in Fiji^78^.

### Centrosomes isolation and fractionation

Centrosomes were isolated from C2C12 cells. Once cells reached 70-80% confluency, they were treated with 1 mM nocodazole and 10 mM Cytochalasin D for 1 h at 37 °C, in order to depolymerize microtubules and actin filaments. After incubation, cells were harvested and resuspended with 8% sucrose in PBS, followed by centrifugation. Cell pellets were resuspended in lysis buffer (8% sucrose, 0.2 mM MgCl_2_ in PBS) and incubated for 5 min on ice, followed by centrifugation for 10 min at 3,500 rpm to remove swollen nuclei and chromatin aggregates. Supernatant was collected and filtered through a nylon mesh. After filtration, supernatant was incubated with 1 μg/ml DNaseI for 30 min and loaded on 30%-60% sucrose gradient, and ultracentrifuged at 130,000 x g for 3 h. Fractions were isolated and analyzed by Western blot with the following markers: Piezo1, Piezo2, γ-Tubulin, Pericentrin and Snx5.

### Cell synchronization and drug treatment

Cells were synchronized at the G2/M boundary by incubating with the CDK1 inhibitor RO-3306 (RO) (Sigma, 9 μM) for 12 h. About 15, 30 and 45 min after drug washout, these cells entered early, mid and late mitotic phase, respectively. For isolating mitotic cells, cells were collected by plate shake off at 30 min after RO washout. For synchronization to the metaphase, cells were treated with 9 μM RO for 12 h, and 25 μM MG132 was added for 1 h after RO washout. For synchronization to G1, cells were incubated with serum starved media (DMEM supplemented 0.1% FBS) for 24 h. For synchronization to S, double thymidine block was used in which 2 mM thymidine was added for 12 h followed by 10 h recovery in regular media, and 2 mM thymidine was added again for 12 h. After thymidine washout, cells were at the G1/S boundary and reached S phase after an additional 3 h in regular media. For Piezo1 and 2 activation or inactivation, cells were incubated with one or a combination of the following compounds: 10 μM Yoda1 (R&D), 10 μM Dooku1 (Glixx Laboratories), 5 μM GsMTx4 (Abcam), 200 μM Jedi1 (Sigma), 200 μM Jedi2 (R&D), 1-20 μM ionomycin (Sigma-Aldrich), 10-40 μM BAPTA-AM (Abcam). For PLK1 inhibition, C2C12 cells were treated with 100 nM BI-6727 (Volasertib) (Selleck Chemicals) for 18 h followed by 30 min treatment with 0.1% DMSO, 10 μM Yoda1 or 5 μM GsMTx4. For microtubules disruption, we treated C2C12 cells for 2 h with 10 μM Taxol (Fisher Scientific), 10 μM Parthenolide (Sigma-Aldrich) or 3h of cold treatment.

### Yeast two-hybrid screens

A yeast two-hybrid (Y2H) screen was performed by Hybrigenics (Paris, France) with the Piezo2 CTD (2683-2752 aa) as the bait on a mixed adult and fetal skeletal muscle library using the fusion vector N-Gal4-Piezo2-CTD. Snx5 was identified as one of the six very high confidence hits (Extended Data Fig. 9b). Validation of the Y2H result was performed by the DupLEX-ATM yeast two-hybrid system (OriGene). Human Piezo2 CTD was cloned into the pEG202NLS vector and fused with LexA. Snx5 and its truncated forms were cloned into the pJG4-5 vector and fused to the transcription activation domain of B42. The lacZ gene in the construct pSH18-34 and the LEU2 gene in the EGY48 strain yeast genome were used as reporters. The pEG202NLS, pSH18-34, and pJG4-5 constructs were co-transformed into yeast EGY48 cells. Interactions were considered positive if yeast cells turned blue in the presence of X-gal and if they grew in the absence of leucine.

### Transmission electron microscopy (EM)

C2C12 cells grown on Aclar plastic film were incubated with 10 μM Yoda1 or 0.1% DMSO (vehicle control) for 1 h. After incubation, cells were fixed by using 2.5% glutaraldehyde, 1.25% PFA and 0.03% picric acid in 0.1 M sodium cacodylate buffer (pH 7.4) for 1 h. Following fixation, cells were washed in 0.1 M sodium cacodylate buffer (pH 7.4), incubated for 30 min in 1% osmium tetroxide (OsO_4_) and 1.5% potassium ferrocyanide (KFeCN6), washed 2x in water and 1x in maleate buffer, incubated in 1% uranyl acetate in maleate buffer for 30 min, washed again twice with water, and subsequently dehydrated in grades of alcohol (5 min each of 50%, 70%, 95%, 2x 100%). Samples were subsequently embedded in TAAB Epon (Marivac Canada Inc., St. Laurent, Canada) and polymerized at 60 °C for 48 h. After polymerization the Aclar film was peeled off and Ultrathin sections (∼80 nm) were cut on a Reichert Ultracut-S microtome, and picked onto copper grids stained with lead citrate. Cells were imaged by using Tecnai G^2^ Spirit BioTWIN at Harvard Medical School EM facility operating at 80 keV with AMT 2k CCD camera.

### Scanning electron microscopy (SEM)

C2C12 cells cultured on Aclar plastic and treated with 10 μM Yoda1, 200 μM Jedi2 or 0.1% DMSO (vehicle control) for 1 h. After incubation, cells were fixed using 2.5% glutaraldehyde, 1.25% PFA and 0.03% picric acid in 0.1 M sodium cacodylate buffer (pH 7.4) for 1 h. Following fixation, cells were washed in 0.1 M sodium cacodylate buffer (pH 7.4), and incubated for 1-2 h in 1% OsO_4_. Next, specimens were dehydrated according to following steps: 50% ethanol for 5 min, 70% ethanol for 5 min, 95% ethanol for 10 min, 100% ethanol for 10 min, and 100% ethanol for 10 min. After dehydration the specimens went through critical point drying for 45-60 min. Specimens were mounted onto stubs and dried overnight. For the final step, specimens were sputter coated with platinum with thickness of 4 nm and imaged with an SEM microscope at Harvard Medical School EM facility.

## Quantification and statistical analysis

Quantitative analysis for all IF experiments were performed based on n=3 independent experiments. Statistical significance was assessed by 2-tailed t-test. ***, ** and * denote p < 0.0001, 0.001 and 0.01, respectively.

---

**Extended Data Fig. 1.**
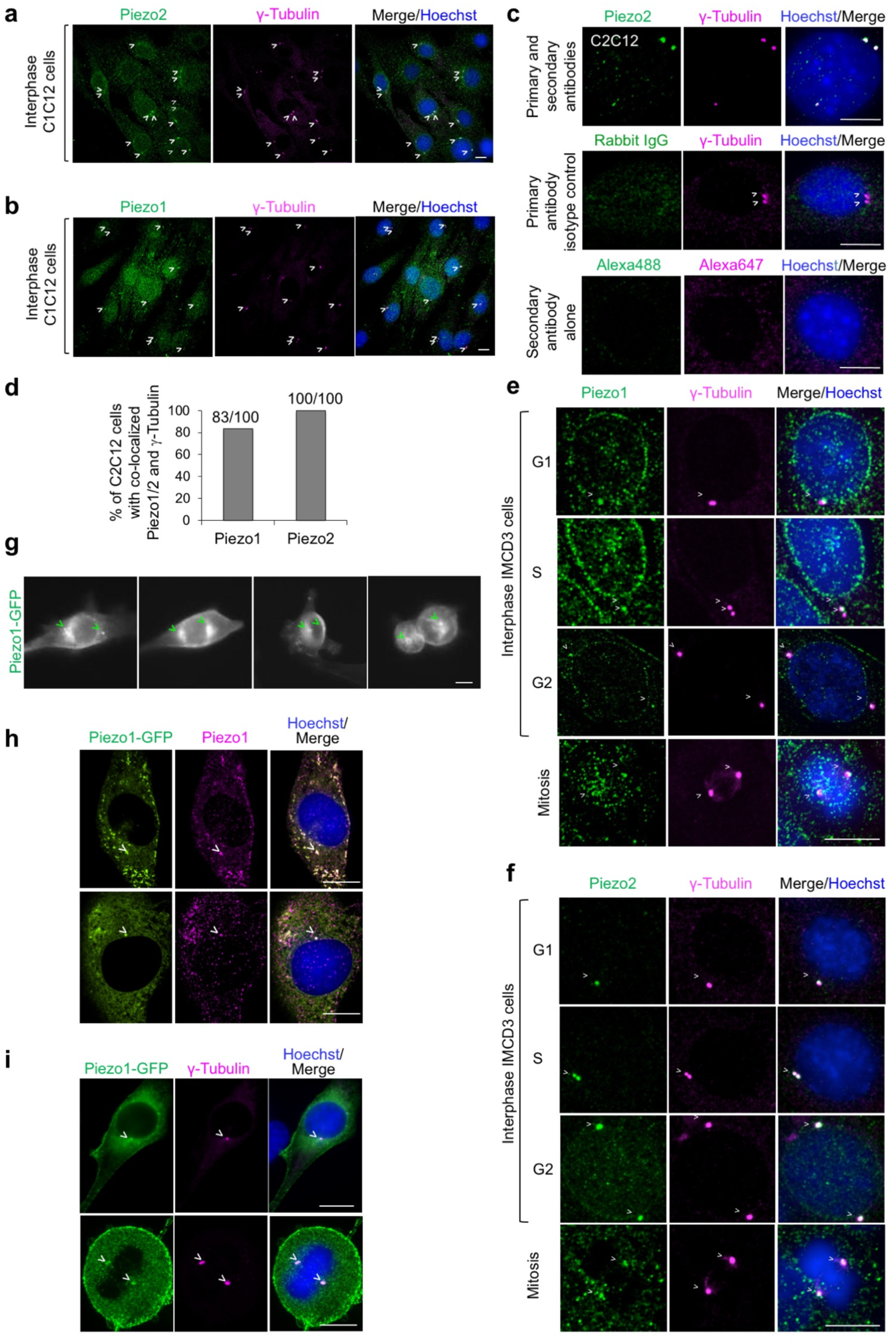
Localization of Piezo1 and 2 at the centrosomes in C2C12 and IMCD3 cells. **a, b,** A field view of interphase C2C12 cells, showing Piezo2 (a) and Piezo 1 (b) localization at the centrosomes. Cells were visualized by IF for Piezo2 (green), γ-Tubulin (magenta), and DNA (Hoechst dye, blue). **c,** Controls for the fixation method, and primary and secondary antibodies in IF. Top: C2C12 cells were fixed in PFA (instead of methanol) and stained using primary antibodies rabbit anti-Piezo2 and mouse anti-γ-Tubulin, and secondary antibodies goat anti-rabbit Alexa Fluor 488 (green) and goat anti-mouse Alexa Fluor 647 (magenta), as well as Hoechst dye for DNA (blue). Middle: C2C12 cells were stained similarly but with a rabbit IgG replacing rabbit anti-Piezo2. Bottom: C2C12 cells were stained only with secondary antibodies. **d,** Quantitative analysis of the percentage of C2C12 cells with co-localized centrosome and Piezo1 or 2 from IF images as in (a) and (b). **e, f,** Centrosome localization of Piezo1 (e) and Piezo2 (f) in IMCD3 cells visualized by IF at different cell cycle stages, imaged by IF as in (a) and (b). **g,** Snapshots from live imaging of C2C12 cells stably expressing Piezo1-GFP during the cell cycle (Supplementary Video 1). Centrosomes are marked with green arrowheads. **h**, Co-localization of Piezo1-GFP fluorescence (green) with anti-Piezo1 IF (magenta). Cells were also stained for DNA (Hoechst, blue). **i,** Localization of Piezo1-GFP fluorescence (green) at the γ-Tubulin (magenta)-marked centrosome by IF. All IF mages are maximum intensity Z projections. Centrosomes and centrosome-localized Piezo1 are marked with white arrowheads. All scale bars are 10 μm, except that the scale bar for (g) is 20 μm.

**Extended Data Fig. 2.**
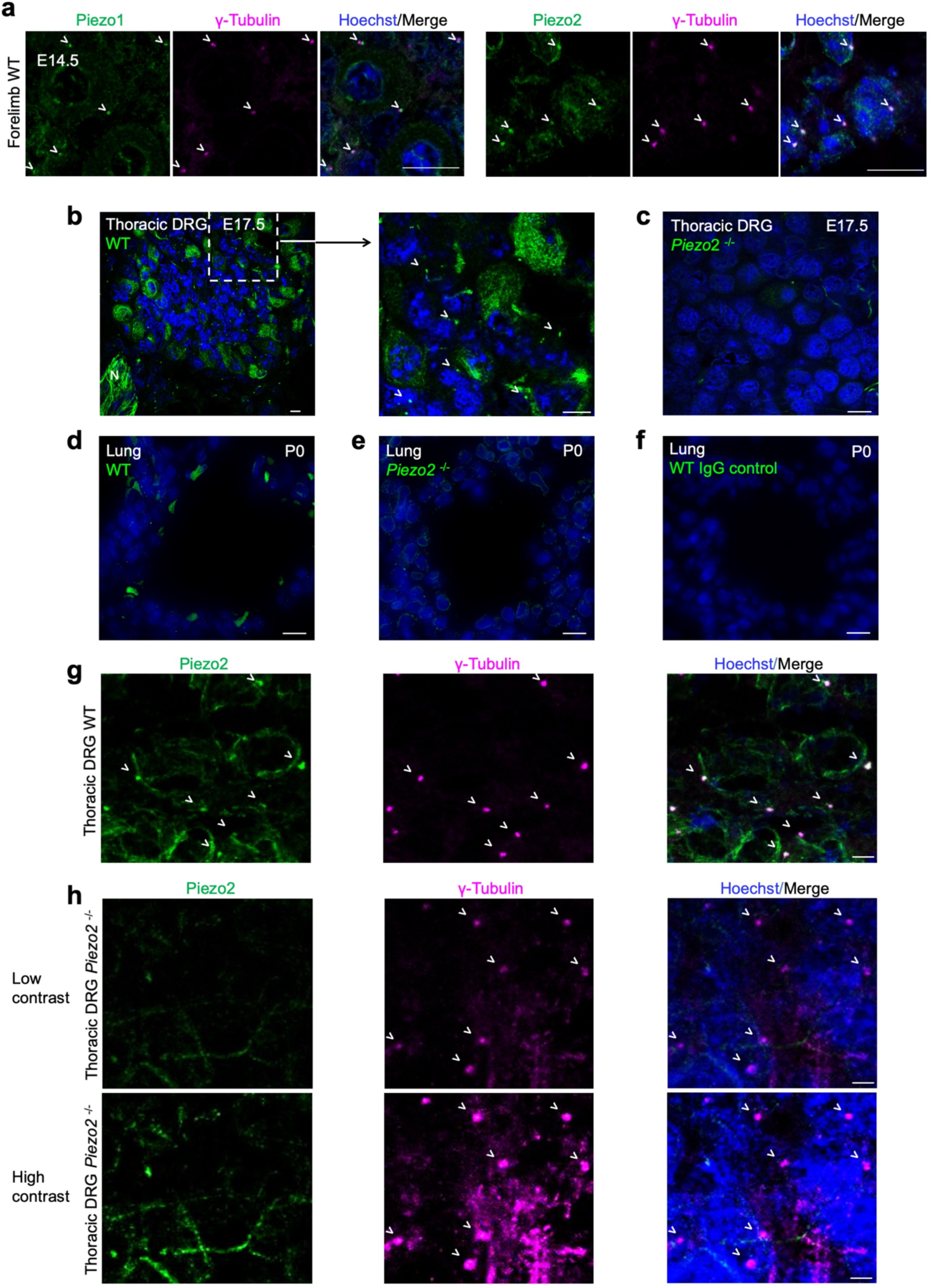
Validation of Piezo2 centrosomal localization by *Piezo2*^-/-^ mice. **a,** Centrosomal localization of Piezo1 (*left)* and Piezo2 (*right*) in E14.5 WT mouse forelimb sections, imaged by IF for Piezo1/2 (green), γ-Tubulin (magenta), and DNA (Hoechst dye, blue). **b,** A thoracic dorsal root ganglion (DRG) section of a WT mouse at E17.5, stained with anti-Piezo2 antibody (green) and Hoechst dye (blue). Right panel shows a 3x zoom-in of the dotted region in the left panel. Piezo2 puncta consistent with a centrosomal localization are labeled with white arrowheads. Of note, Piezo2 positive puncta are often present in DRG neurons whether or not the cell bodies are also Piezo2 positive. **c,** A thoracic DRG section of a *Piezo2*^-/-^ KO mouse at E17.5, stained with anti-Piezo2 antibody (green) and Hoechst dye (blue). Piezo2 expression at DRG neuronal cell body or centrosome is largely undetectable**. d,** A lung bronchial section of a WT mouse at P0, stained with anti-Piezo2 antibody (green) and Hoechst dye (blue). The cells labeled in green represent scattered pulmonary neuroepithelial cell bodies (NEBs). **e,** A comparable lung bronchial section of a *Piezo2*^-/-^ KO mouse at P0, imaged by IF with anti-Piezo2 antibody and Hoechst dye (blue), showing absence of Piezo2 staining. **f,** A comparable lung bronchial section of a WT mouse at P0, stained with rabbit IgG (green) as primary antibody and Hoechst dye (blue). **g, h,** Thoracic DRG sections of WT mice (g) and *Piezo2*^-/-^ KO mice, shown at 2 different contrast levels (h) at E17.5, stained with anti-Piezo2 antibody (green), γ-Tubulin (magenta) and Hoechst dye (blue). White arrowheads identify γ-Tubulin-marked centrosomes (magenta) and/or Piezo2. All scale bars are 10 μm.

**Extended Data Fig. 3.**
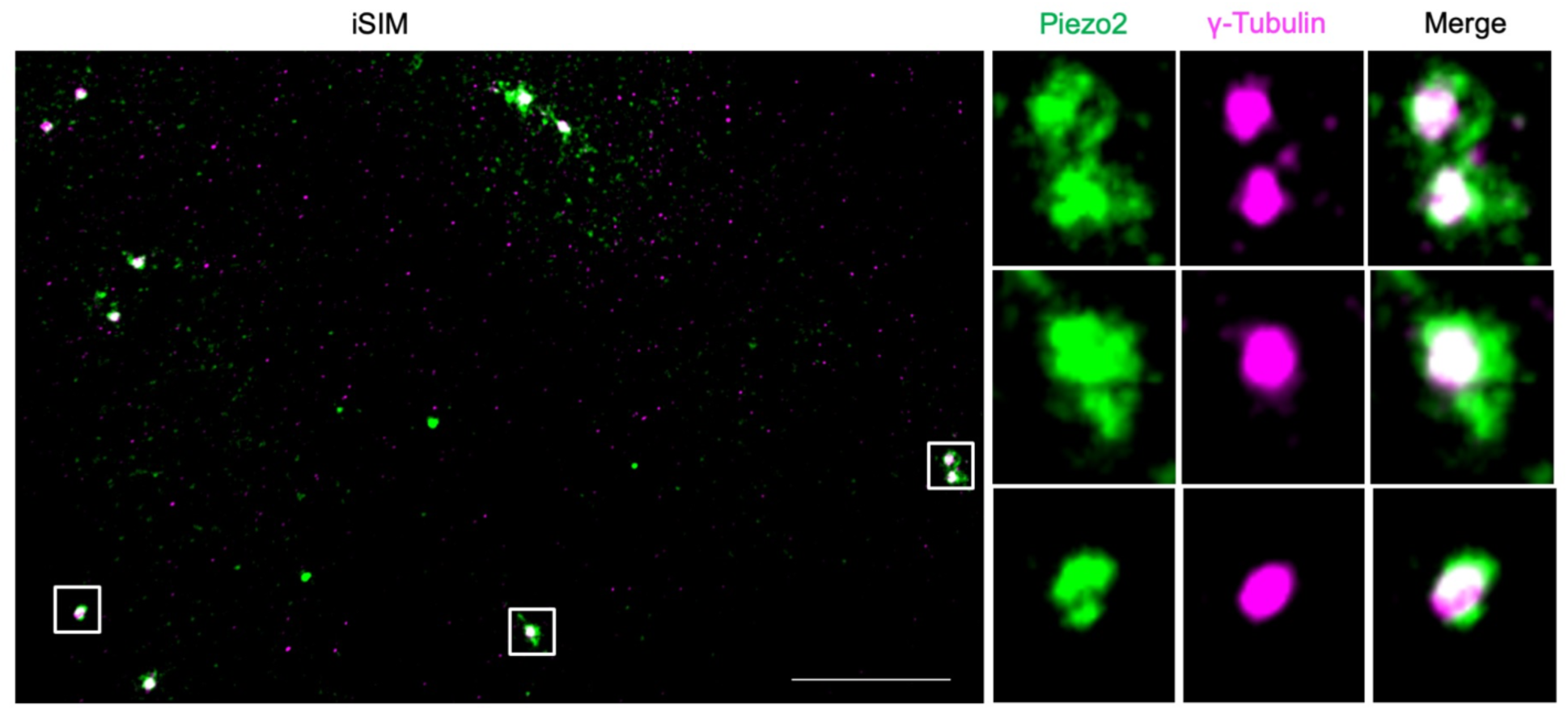
Instant structured illumination microscopy (iSIM) for Piezo2 and γ-Tubulin in C2C12 cells. iSIM was performed for fixed unsynchronized C2C12 cells, stained with Piezo2 (green) and γ-Tubulin (magenta). The iSIM image (left) shows co-localization of Piezo2 and γ-Tubulin, and the insets of the centrosomes (right) are shown with x4 zoom-in.

**Extended Data Fig. 4.**
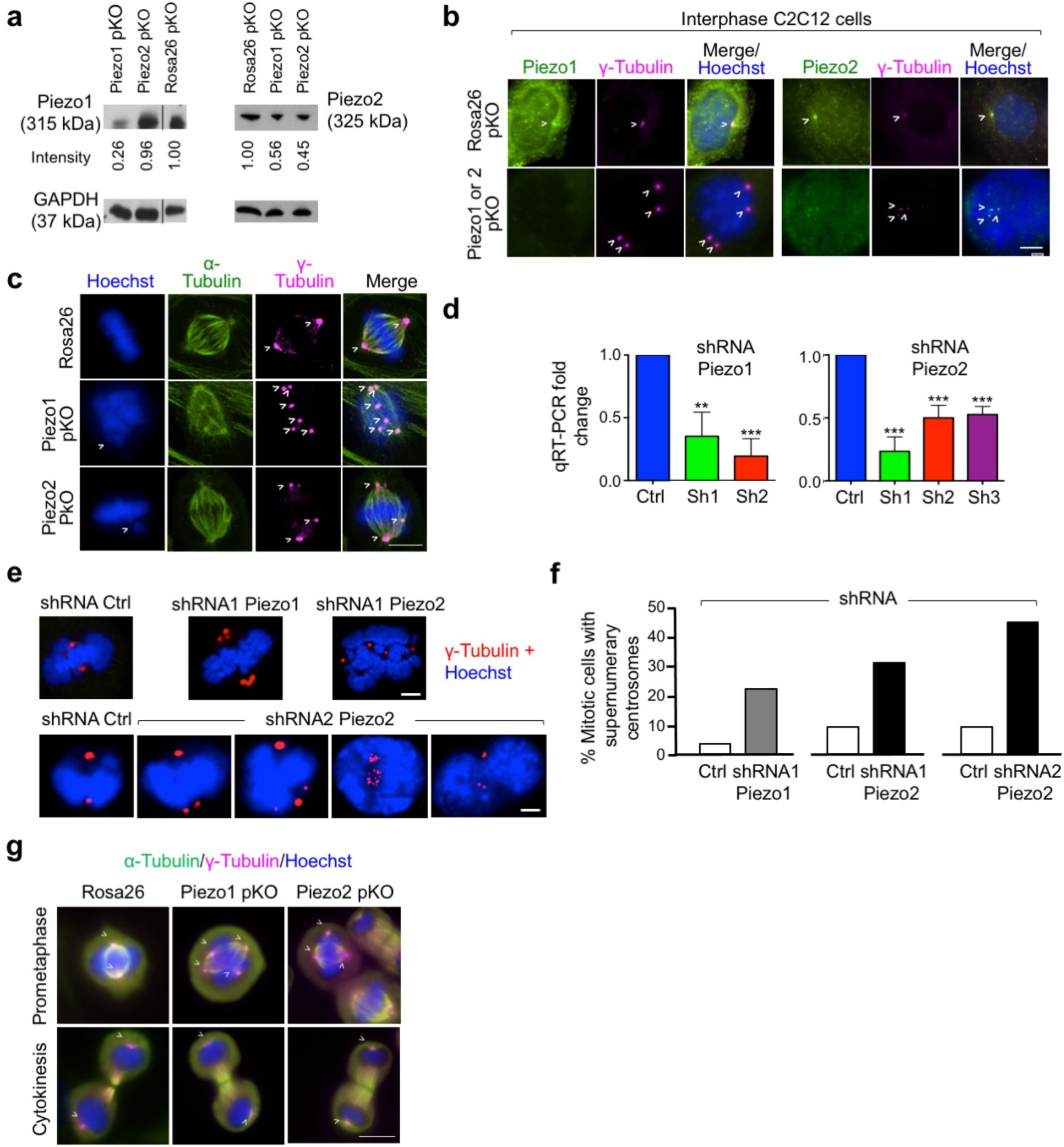
Piezo1 and 2 CRISPR-Cas9 pKO and ShRNA KD in C2C12 cells. **a,** Piezo1 and Piezo2 protein levels in Piezo1 pKO, Piezo2 pKO, or Rosa26 pKO (control) of C2C12 cells analyzed by Western blots with GAPDH as the loading control. Densitometric analysis of Western blot bands of Piezo1 or 2 are shown. **b,** Piezo1 pKO (*left*) and Piezo2 pKO (*right*) interphase C2C12 cells at day 1 post-pKO selection, imaged by IF for Piezo1 or Piezo2 (green), γ-Tubulin (magenta) and DNA (Hoechst, blue). Centrosomes are marked with white arrows, and supernumerary centrosomes are seen in Piezo pKO cells, in comparison with Rosa26 pKO control. **c,** Piezo1 pKO (*middle row*), Piezo2 pKO (*bottom row*) and Rosa26 pKO (*top row*) of mitotic C2C12 cells at day 1 post-pKO selection, imaged by IF for α-Tubulin (green), γ-Tubulin (magenta) and DNA (Hoechst, blue). Centrosomes and lagging chromatin are marked with white arrows, and misaligned microtubules are seen in Piezo pKO cells, in comparison with the Rosa26 pKO control **d,** Quantitative real-time RT-PCR (qRT-PCR) of Piezo 1 or 2 transcripts upon small hairpin RNA (shRNA) KD of Piezo1 or Piezo2 in C2C12 cells using two and three different shRNAs for Piezo1 and Piezo2, respectively, in comparison with an off target shRNA against firefly luciferase mRNA as the control. **e,** Control KD, Piezo1 shRNA KD (one shRNA) or Piezo2 shRNA KD (two shRNAs) C2C12 cells at day 4 post-KD selection. The cells were imaged by IF for γ-Tubulin (red) and DNA (Hoechst, blue), showing supernumerary centrosomes upon Piezo1 or Piezo2 KD. **f,** Quantitative analysis of C2C12 cells with supernumerary centrosomes upon Piezo1 shRNA KD (one shRNA) or Piezo2 shRNA KD (two shRNAs) at day 4 post-KD selection. All scale bars are 10 μm. **g,** Off target, Piezo1 and 2 pKO C2C12 cells at day 1 post selection and 45 min after release of G2/M border synchronization with RO-3066. Cells were imaged by IF for α-tubulin (green), γ-tubulin (magenta) and DNA (Hoechst dye, blue), shown at prometaphase and cytokinesis. Quantitative analysis of mitotic cell populations from the IF experiments shown in Fig. 1i. Statistical significance between an experimental group and the control group was assessed by 2-tailed t-test. ***, ** and * denote p < 0.0001, 0.001 and 0.01, respectively.

**Extended Data Fig. 5.**
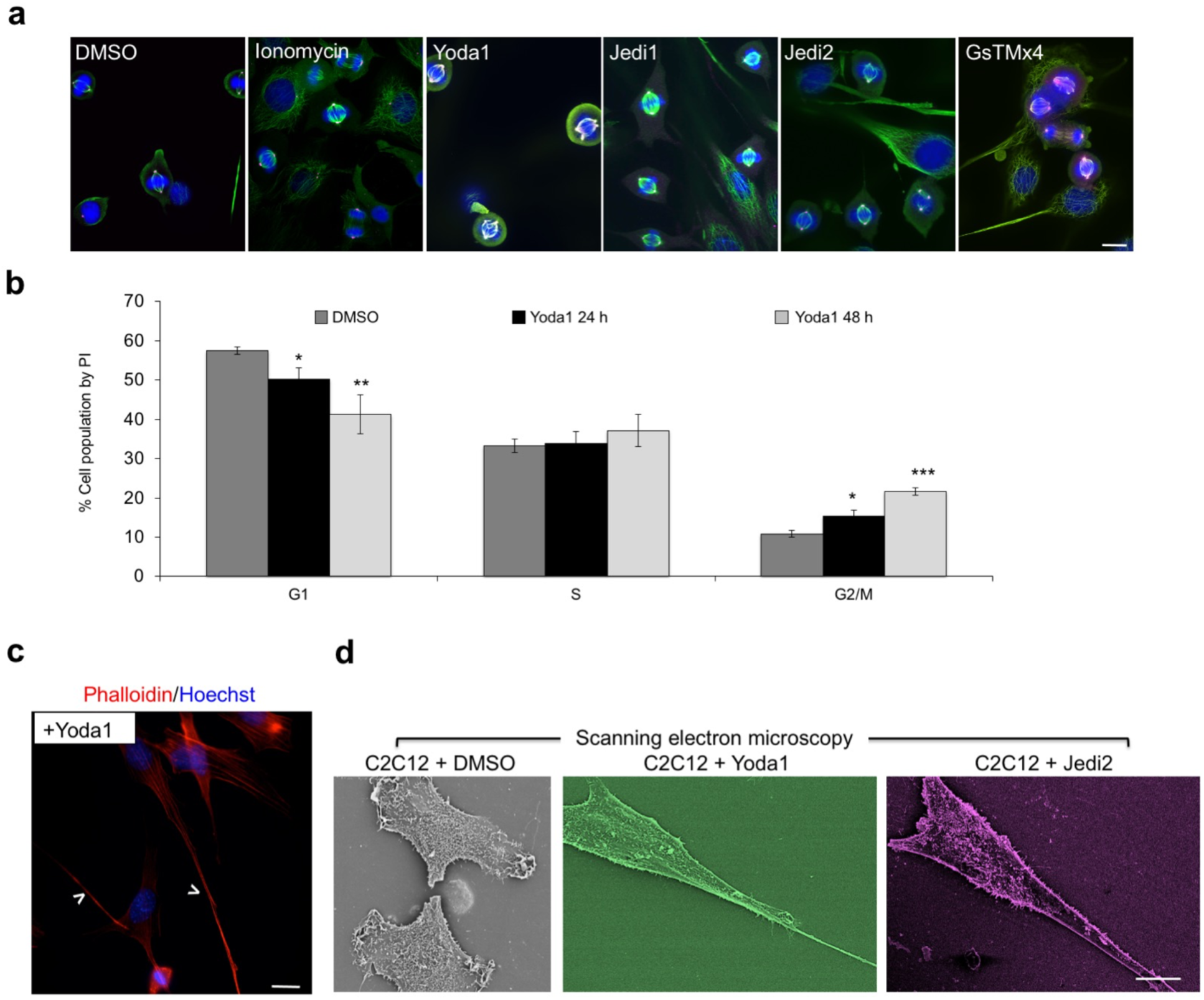
Intracellular and cell surface effects upon Piezo activation and inhibition. **a,** Representative field views of C2C12 cells 30 min after drug treatment visualized by IF for α-Tubulin (green), γ-Tubulin (magenta) and DNA (Hoechst, blue). Drugs – DMSO alone (0.1%, vehicle control), Yoda1 (10 μM), GsMTx4 (5 μM), Jedi1 (200 μM), Jedi2 (200 μM), or ionomycin (20 μM) – were added upon release of G2/M synchronization by RO-3306. Images are maximum intensity Z projections. **b,** Cell cycle effect of Piezo1 activation 24 and 48 h after Yoda1 addition. The cells were analyzed using PI (propidium iodide) staining and flow cytometry, showing increasing accumulation in G2/M with longer Yoda1 treatment. Statistics represent 3 independent experiments, analyzed by two-tailed t-test with ***, ** and * for p < 0.0001, 0.001 and 0.01, respectively. **c,** C2C12 cells treated with Yoda1 for 1 h imaged by phalloidin-staining of F-actin (red) and Hoechst-staining of DNA (blue). Protrusions are marked with white arrowheads. **d,**Scanning electron microscopy (SEM) images of C2C12 cells after 1 h treatment with Yoda1 (10 μM), Jedi2 (200 μM) or DMSO (0.1 %). All scale bars: 10 μm.

**Extended Data Fig. 6.**
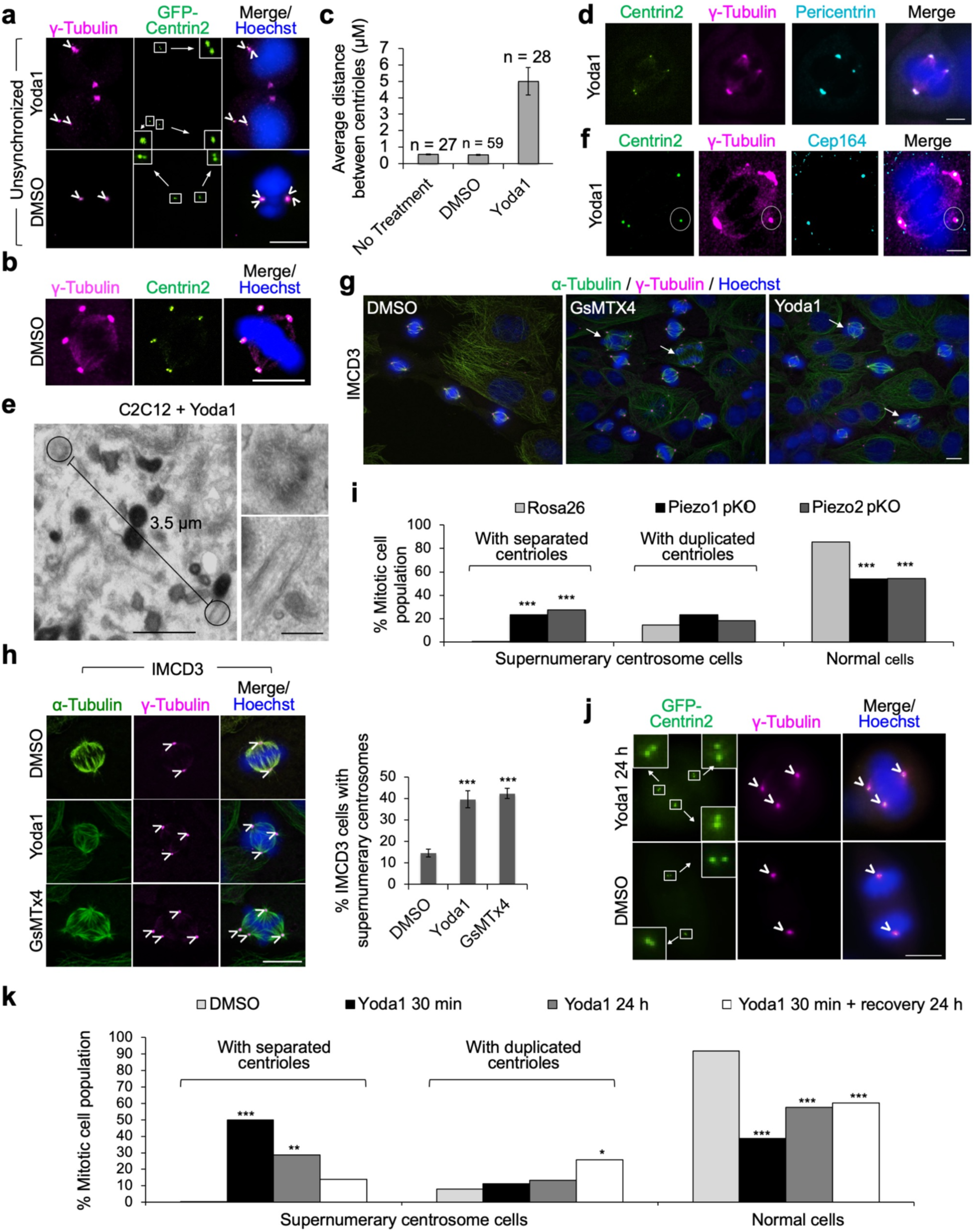
Centriole disengagement upon Piezo protein activation and inhibition. **a,** Unsynchronized C2C12 cells expressing GFP-Centrin2 (green), fixed 30 minutes after introducing Yoda1 (10 μM) or DMSO (0.1%, vehicle control) and stained for γ-Tubulin (magenta) and DNA (Hoechst, blue). Yoda1 treatment resulted in centriole disengagement. **b,** Supernumerary centrosomes in RO-3306-synchronized C2C12 cells expressing GFP-Centrin2 (green) fixed 30 minutes after introducing 0.1% DMSO and stained for γ-Tubulin (magenta) and DNA (Hoechst, blue). The low percentage of supernumerary centrosomes in DMSO control was resulted from centriole duplication**. c,** Analysis of the measured disengagement distance between centrioles upon Piezo1 activation using IF experiments as in (a). **d,** C2C12 cells expressing GFP-Centrin2 (green), fixed 30 minutes after introducing Yoda1 (10 μM) and stained for γ-Tubulin (magenta), Pericentrin (cyan) and DNA (Hoechst, blue), confirming the centrosomal defect. **e,** A negative-stain EM image of separated centrioles in a thin plastic section of embedded C2C12 cells 30 min after Yoda1 treatment (left) and its zoom-ins (right). Two centrioles are in a cross-sectional view with the normal 9-fold symmetry and a longitudinal view, respectively, at a disengagement distance of 3.5 μm. **f,** C2C12 cells expressing GFP-Centrin2 (green), fixed 30 minutes after introducing Yoda1 (10 μM) and stained for γ-Tubulin (magenta), Cep164 (cyan) and DNA (Hoechst, blue). Cep164 marks mother centrioles, and upon Yoda1 addition, the daughter centriole specifically separated from the mitotic spindle, shown in a white circle. **g,** Fields of view of G2/M synchronized IMCD3 cells 30 min after treatment with DMSO (0.1%), Yoda1 (10 μM) or GsMTx4 (5 μM), imaged by IF for α-Tubulin (green), γ-Tubulin (magenta) and DNA (blue). Synchronization was done using RO-3306 and drugs were added after RO-3306 washout. Cells containing supernumerary centrosomes are marked with white arrows. **h,** Detailed views of IMCD3 cells treated as in (g) (*left*) and quantitative analysis of cells with supernumerary centrosomes (*right*). **i,** Quantitative analysis of mitotic Piezo1 and 2 pKO cells at day 1 post-KO selection with supernumerary centrosomes that contain either separated or duplicated centrioles visualized by IF. Among cells with supernumerary centrosomes, significantly more of them contained separated centrioles in Piezo pKO cells compared to Rosa26 control cells. **j**, Supernumerary centrosomes with duplicated centrioles in GFP-Centrin2 C2C12 cells after 24 h treatment with Yoda1 (10 μM), imaged for GFP-Centrin2 fluorescence (green), γ-Tubulin IF (magenta) and DNA (blue). **k,** Quantitative analysis of mitotic cells with supernumerary centrosomes that contain either separated or duplicated centrioles upon different Yoda1 treatments, visualized by IF as in (j). The percentage of supernumerary centrosome cells with duplicated centrioles is higher after a 24 h recovery period post the 30 min Yoda1 treatment in comparison with either 30 min or 24 h Yoda1 treatment alone. By contrast, the percentage of supernumerary centrosome cells with separated centrioles is lower after a 24 h recovery period post the 30 min Yoda1 treatment. Scale bars are 10 μm for all IF images, 1 μm for the left panel in (e), and 200 nm for the right zoom-ins in (e). All IF mages are maximum intensity Z projections and data are represented by three independently quantified experiments counting 50-250 cells each. Statistical significance between an experimental group and the control was assessed by two-tailed t-test with ***, ** and * for p < 0.0001, 0.001 and 0.01, respectively.

**Extended Data Fig. 7.**
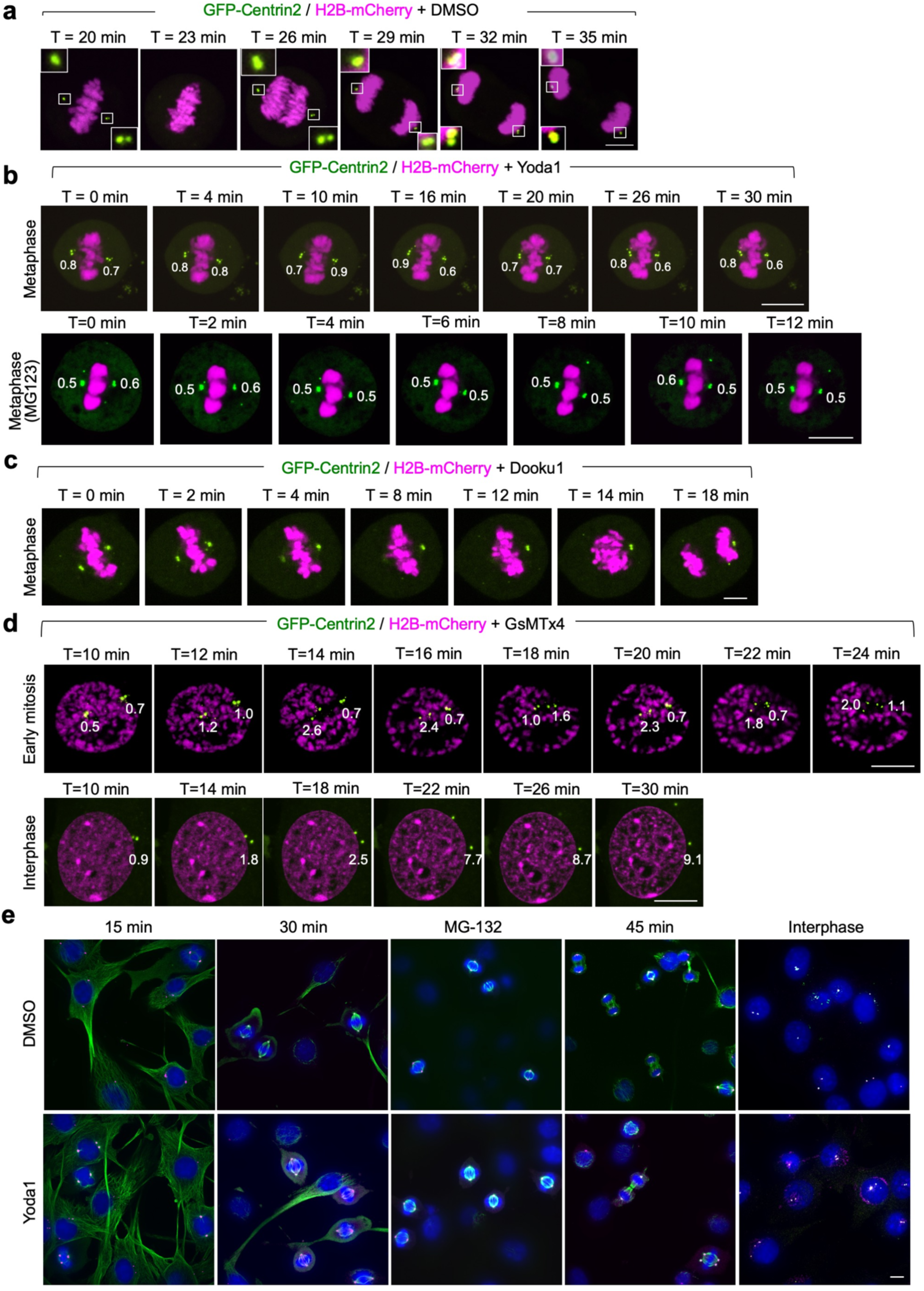
Centriole disengagement upon Piezo1 activation or Piezo inhibition in live and fixed cells. **a-d,** Movie snapshots of imaged C2C12 cells stably expressing GFP-Centrin2 (green) and H2B-mCherry (magenta) treated with DMSO (0.1%, vehicle control) (a), Yoda1 (10 μM) at metaphase 30 minutes after RO-3306 release (b, top) or after MG132 synchronization (b, bottom), Dooku1 (10 μM) at metaphase or GsMTx4 (5 μM) at both early mitosis and interphase (d). Distances between centrioles are labeled in μm. The DMSO-treated cell contained centrosomes with two centrioles separated by < 0.6 μm. The Yoda1 or Dooku1-treated cell at metaphase did not show separated centrioles. The GsMTx4-treated cells at early mitosis and interphase both exhibited centriole disengagement. **e**, Representative field views of C2C12 cells treated with DMSO (top row) or Yoda1 (bottom row) for 10 min at 15 min (prophase and prometaphase), 30 min (metaphase) or 45 min (telophase and cytokinesis) after RO-3306 release, after MG132 synchronization (metaphase) or with double thymidine block (interphase). Cells were imaged by IF for α-Tubulin (green), γ-Tubulin (magenta) and DNA (blue). Centriole disengagement was observed in interphase, prophase and prometaphase, and telophase and cytokinesis, but not in metaphase. All images are maximum intensity Z projections and all scale bars are 10 μm.

**Extended Data Fig. 8.**
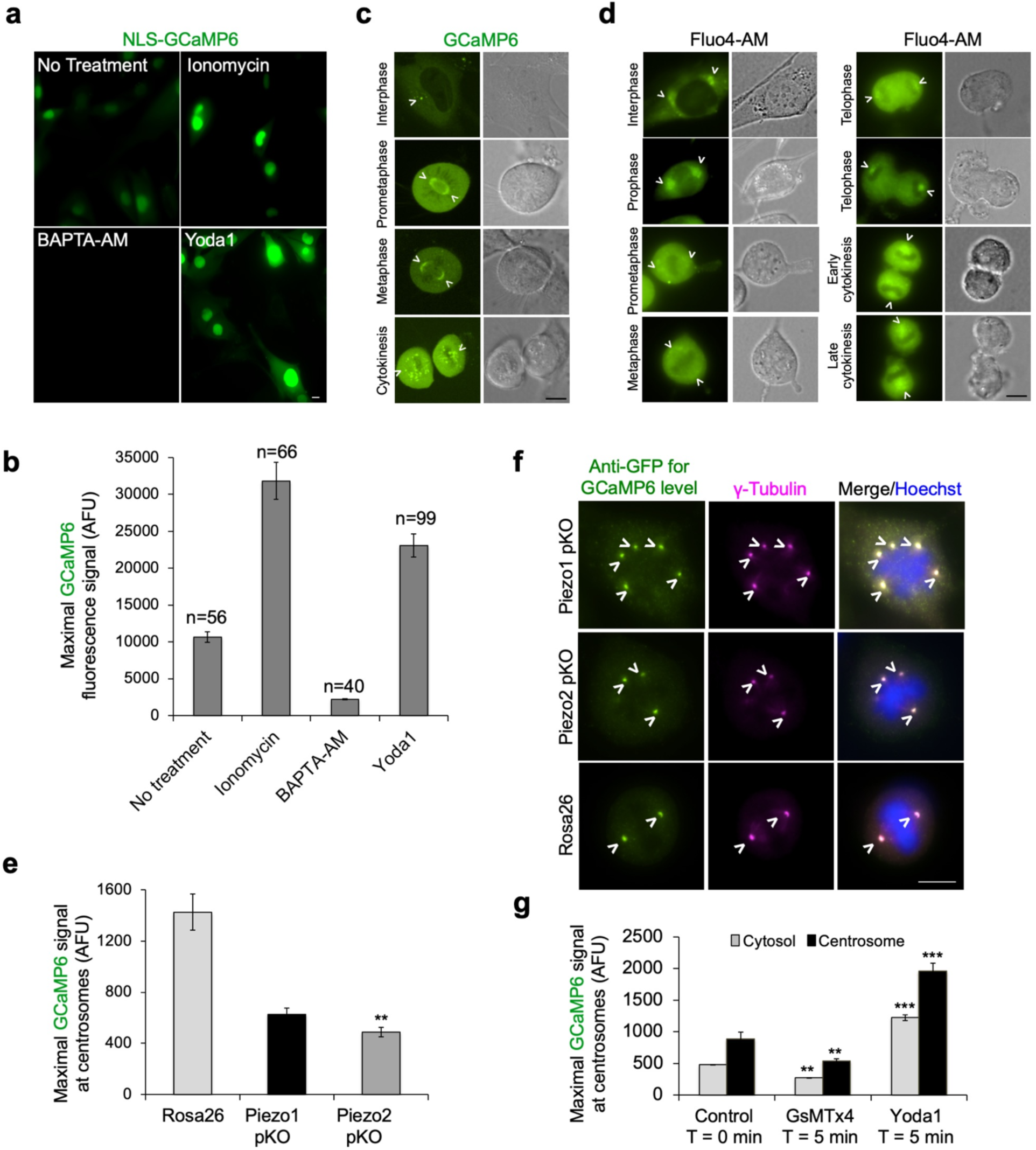
Calcium imaging by the Ca^2+^-sensitive GCaMP6 reporter. **a,** Validation of the NLS-GCaMP6 reporter stably expressed in C2C12 cells by ionomycin (1 μM), BAPTA-AM (10 μM), or Yoda1 (10 μM) treatment. **b**, Quantitative analysis of maximal signals for cells imaged in (a). AFU denotes arbitrary fluorescence units. **c,** Phase and fluorescence snapshots of C2C12 cells expressing GCaMP6 reporter at interphase and different stages of mitosis (green). Concentrated Ca^2+^ signals that represent centrosome locations are marked by white arrowheads. **d**, Phase and fluorescence snapshots of C2C12 cells at interphase and different stages of mitosis, imaged live at 45 min after treatment with the Ca^2+^ dye Fluo4-AM (9 μM, green). Concentrated Ca^2+^ signals that represent centrosome locations are marked by white arrowheads. **e,** Quantitative analysis of maximal GCaMP6 signals at centrosomes as shown in Fig. 4b, revealing reductions to 43% and 34% in comparison to Rosa26 pKO control values in Piezo1 and 2 pKO cells, respectively. **f,**GCaMP6 expression by anti-GFP IF (green) in NLS-GCaMP6 expressing C2C12 cells at day 1 post-KO selection of Rosa26 pKO, Piezo1 pKO or Piezo2 pKO. IF for γ-Tubulin (magenta) and Hoechst dye staining of DNA (blue) are also shown, exhibiting equal GCaMP6 expression regardless of Piezo1 or 2 pKO. **g**, Quantitative analysis of maximal Ca^2+^ signal indicated by GCaMP6 fluorescence intensity at centrosomes and in the cytosol as shown in Fig. 4c, showing that there was a similar increase or decrease of local Ca^2+^ concentration upon Yoda1 activation or GsMTx4 inhibition relative to the untreated control value both in the cytosol and at the centrosomes. Images are maximum intensity Z projections and all scale bars are 10 μm. Data are represented by mean ± SEM from three independently quantified experiments. Statistical significance between an experimental group and the control was assessed by 2-tailed t-test with *** and ** for p < 0.0001 and 0.001, respectively.

**Extended Data Fig. 9.**
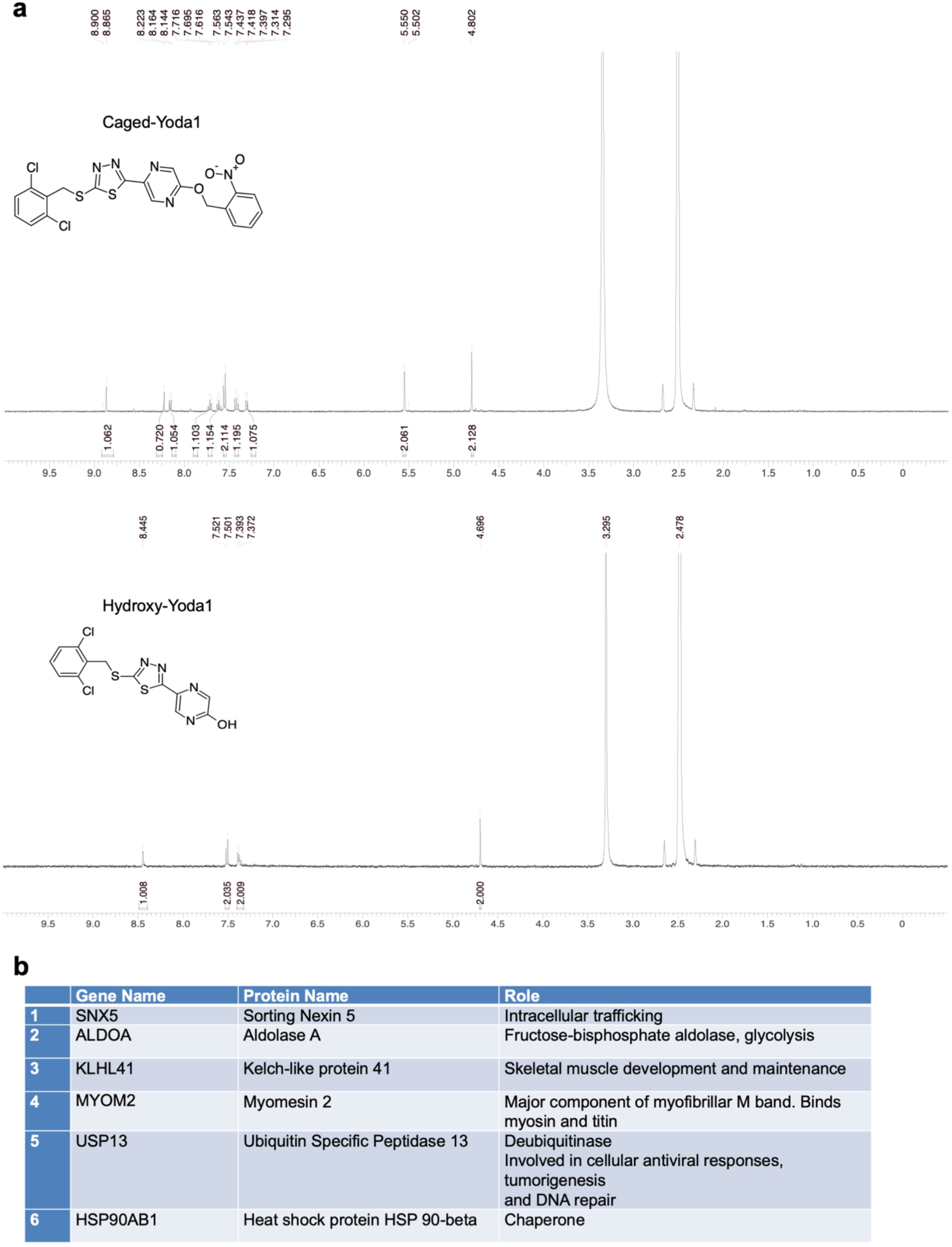
Chemical synthesis quality control and yeast two-hybrid screen result. **a,** HNMR Spectra of caged-Yoda1 and hydroxy-Yoda1 synthesis. The compounds were synthesized by WuXi AppTec with purity determined by LC-MS of 98.2% and 78.6%, respectively. For hydroxy-Yoda1, the compound was clean based on the HNMR spectrum but might not be stable in LC-MS and therefore a lower purity was reported. **b,** A table describing the top hits from the yeast two-hybrid screen performed for Piezo2 CTD.

**Extended Data Fig. 10.**
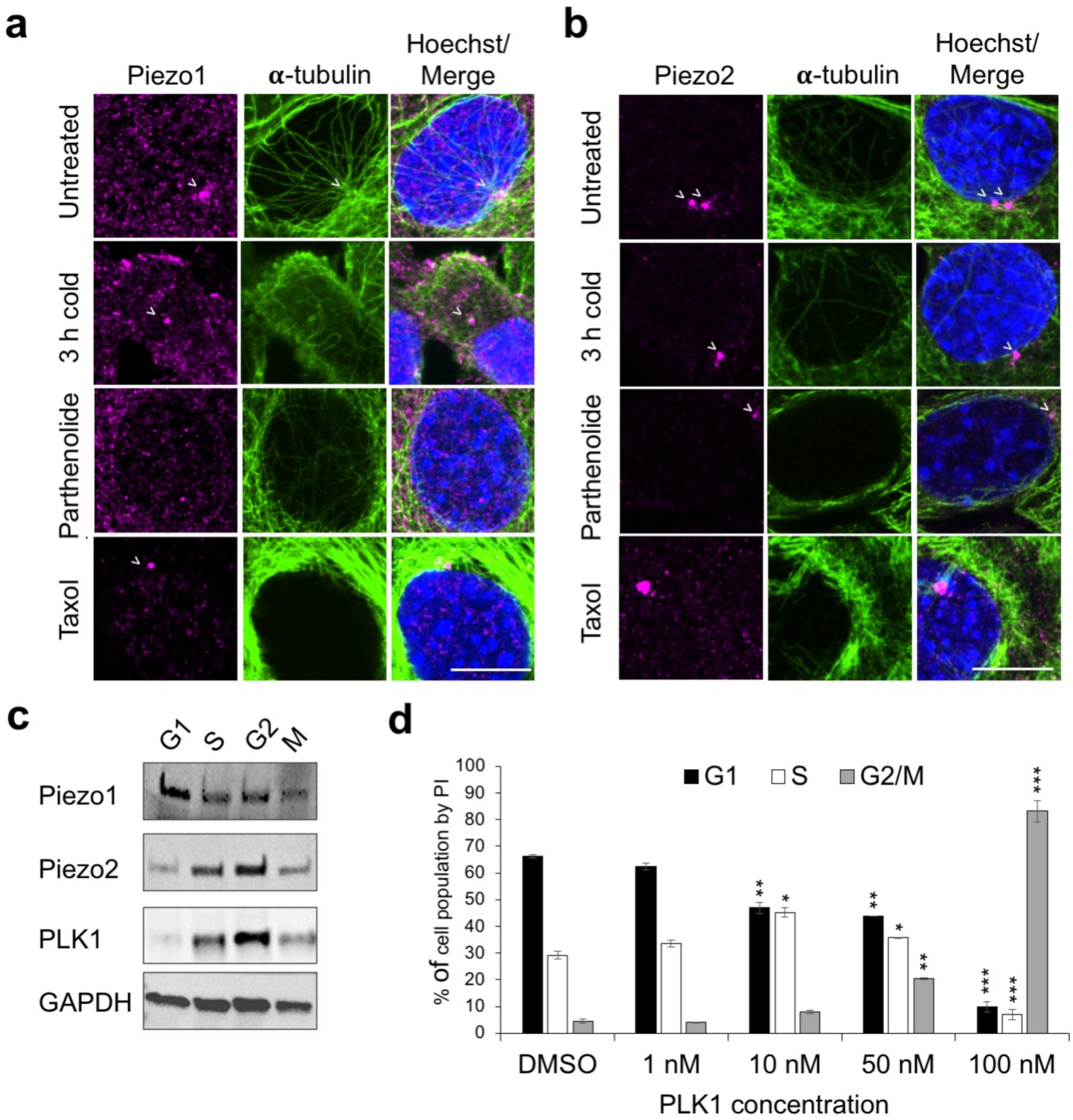
Microtubule disruption compromised Piezo1/2 centrosomal localization. **a-b,** IF of C2C12 cells, fixed after 3h cold treatment, 2 h treatment with Parthenolide or Taxol and stained with Piezo1 (a) or Piezo2 (b), ⍺-Tubulin (green) and Hoechst (Blue). Untreated cells were used as a control. Centrosomes are marked with white arrowheads. **c**, Western blots of C2C12 cells at different cell cycle stages using anti-Piezo1, Piezo2, PLK1 and GAPDH (loading control) antibodies. While Piezo2 is highly expressed at S and G2, Piezo1 expression is relatively constant. **d**, Cell cycle analysis by PI staining for C2C12 cells treated with PLK1 inhibitor BI-6727. Optimal concentration for BI-6727 treatment in C2C12 was determined as 100 nM. Images are maximum intensity Z projections and all scale bars are 10 μm. Data are represented by mean ± SEM from three independently quantified experiments. Statistical significance between an experimental group and the control was assessed by 2-tailed t-test with ***, ** and * for p < 0.0001, 0.001 and 0.01, respectively.

**Supplementary Video 1. Live cell imaging of a C2C12 cell stably expressing Piezo1-GFP**. Cells were imaged on a confocal microscope using a 20x objective with 1.5 zoom for 5.5 h with 5 min intervals.

**Supplementary Video 2**. **Live cell imaging of a C2C12 cell stably expressing GFP-Centrin2 and H2B-mCherry following treatment with DMSO (0.1%), showing normal cell division**. Cells were imaged for 24 min at 3 min intervals using a 60x objective on a spinning disk confocal microscope 20 min after RO release.

**Supplementary Video 3-8**. **Live imaged C2C12 cells stably expressing GFP-Centrin2 and H2B-mCherry upon Yoda1 (10 μM) addition.** Supplementary Video 3 captured a prophase cell 15 min after RO-3306 release, showing centriole disengagement from intact centrosomes and the appearance of lagging chromatin upon Yoda1 addition. The video is 30 min in length over 30 frames (1 min per frame). Supplementary Video 4 imaged a metaphase cell 30 min after RO-3306 release upon Yoda1 addition. The video is 30 min in length over 15 frames (2 min per frame). Supplementary Video 5 captured a cell at cytokinesis 45 min after RO-3306 release, showing centriole disengagement upon Yoda1 addition. The length of the video is 12 min over 4 frames (3 min per frame). Supplementary Video 6 imaged unsynchronized interphase cell at G1, showing centriole disengagement upon Yoda1 addition. The length of the video is 30 min with 15 frames (2 min per frame). Supplementary Video 7 imaged unsynchronized interphase cell at G2, showing centriole disengagement upon Yoda1 addition. The length of the video is 20 min with 10 frames (2 min per frame). Supplementary Video 8 imaged a metaphase cell synchronized by MG132 upon Yoda1 addition. The length of the video is 20 min with 10 frames (2 min per frame). Imaging began immediately upon Yoda1 addition (Supplementary Video 3-4, 6-8) or 5 min after Yoda1 addition (Supplementary Video 5) using a 60x objective on a spinning disk confocal microscope.

**Supplementary Video 9**. **Live imaged C2C12 cells stably expressing GFP-Centrin2 and H2B- mCherry upon Dooku1 (10 μM) addition**. Supplementary Video 9 imaged a metaphase cell 30 min after RO-3306 release upon Dooku1 addition. The video is 20 min in length over 10 frames (2 min per frame).

**Supplementary Video 10-11**. **Live imaged C2C12 cells stably expressing GFP-Centrin2 and H2B-mCherry upon GsTMx4 (5 μM) addition**. Supplementary Video 10 documents a prophase cell and Supplementary Video 11 presents an interphase cell, showing centriole disengagement from intact centrosomes upon GsTMx4 addition. Imaging began 10 min after GsTMx4 addition, with 2 min intervals using a 60x objective on a spinning disk confocal microscope. The length of Supplementary Video 10 is 16 min with 8 frames, and length of Supplementary Video 11 is 20 min with 10 frames (2 min per frame).

**Supplementary Video 12**. **Live cell imaging of C2C12 cells stably expressing NLS-GCaMP6 during mitosis.** Cells were imaged for 35 min at 5 min intervals using a 60x objective on a spinning disk confocal microscope.

**Supplementary Video 13-14**. **Live cell imaging of C2C12 cells stably expressing GCaMP6-GTU during interphase.** Cells were imaged for 20 min at 2 min intervals using a 60x objective on a spinning disk confocal microscope upon addition of Yoda1 (Supplementary Video 14) or vehicle only (Supplementary Video 13).

**Supplementary Video 15. Live cell imaging of C2C12 cells stably expressing GFP-Centrin2 upon uncaging of caged-Yoda1 at the centrosome**. Cells were imaged on a confocal microscope after uncaging caged-Yoda1 at the centrosome with a 405 nm laser using a 63x objective for 20 min with an interval of 8 sec. The field of view shows a GFP-Centrin2 cell that was uncaged, in addition to a control GFP-Centrin2 cell that remained caged.

**Supplementary Video 16. Live cell imaging of C2C12 cells stably expressing GFP-Centrin2 upon uncaging of caged-Yoda1 at the centrosome**. The cell was imaged on a confocal microscope after uncaging caged-Yoda1 at the centrosome with a 405 nm laser using a 63x objective for 10 min with an interval of 20 sec. Of note, no cell surface protrusion was observed, supporting local Piezo1 activation at the centrosome.

**Supplementary Video 17. Live cell imaging of C2C12 cells stably expressing GFP-Centrin2 upon uncaging of caged-Yoda1 away from at the centrosome**. The cell was imaged on a confocal microscope after uncaging caged-Yoda1 in a different area of the cell (not at the centrosome) with a 405 nm laser using a 63x objective for 8 min with an interval of 10 sec.

